# Social experience and pheromone receptor activity reprogram behavioral switch and neuromodulatory gene expression in sensory neurons

**DOI:** 10.1101/2021.06.18.449021

**Authors:** Bryson Deanhardt, Qichen Duan, Chengcheng Du, Charles Soeder, Alec Morlote, Deeya Garg, Corbin D. Jones, Pelin Cayirlioglu Volkan

## Abstract

Social experience and pheromone signaling in olfactory neurons affect neuronal responses and male courtship behaviors in *Drosophila*. We previously showed that social experience and pheromone signaling modulate chromatin around behavioral switch gene *fruitless*, which encodes a transcription factor necessary and sufficient for male behaviors. Fruitless drives social experience dependent modulation of courtship behaviors and pheromone responses in sensory neurons, however, the molecular mechanisms underlying this neuromodulation remain less clear. To identify the molecular mechanisms driving social experience-dependent neuromodulation, we performed RNA-seq from antennal samples of mutants in pheromone receptors and *fruitless,* as well as grouped or isolated wild-type males. We found that loss of pheromone detection differentially alters the levels of *fruitless* exons suggesting changes in splicing patterns. In addition, many Fruitless target neuromodulatory genes, such as neurotransmitter receptors, ion channels, ion and membrane transporters, and odorant binding proteins are differentially regulated by social context and pheromone signaling. Recent studies showed that social experience and juvenile hormone signaling coregulate *fru* chromatin to modify pheromone responses in olfactory neurons. Interestingly, genes involved in juvenile hormone metabolism are also misregulated in different social contexts and mutant backgrounds. Our results suggest that modulation of neuronal activity and behaviors in response to social experience and pheromone signaling likely arise due to large-scale changes in transcriptional programs for neuromodulators downstream of behavioral switch gene function.

## Background

Detection of the social environment through pheromone signaling is critical for animals to recalibrate sex-specific behaviors such as mating and aggression [1–4]. It is thought that changes in social environment can modify the regulation of genes necessary for neuronal homeostasis, physiology, and transmission, ultimately affecting circuit function and behaviors [2, 5, 6]. Previous studies on the effects of early life experience have identified changes in neuroanatomy, synaptic plasticity, neurotransmission, and gene expression. For example, maternal licking and grooming of pups, increases DNA methylation around glucocorticoid receptor gene, leading to long-lasting effects on offspring stress responses and behaviors [7–9]. However, transcriptional cascades driving sensory and social experience-dependent modulation of gene expression, circuit function, and behaviors remain unclear.

Identifying gene regulation cascades by which social signals influence neural and behavioral responses requires a model system with well-defined circuits and genetic regulators with roles in neurophysiology, circuit structure, and behavioral function. Circuitry for courtship behavior in *Drosophila melanogaster* is an excellent experimental system to address this question. In *Drosophila*, male-specific courtship behaviors are governed by a critical transcriptional regulator Fruitless^M^ (Fru^M^), which is encoded by the male-specific alternative splicing of the *fruitless* (*fru*) gene from the P1 promoter [10, 11]. It is known that Fru^M^ is both necessary and sufficient for male courtship as loss of Fru^M^ in males leads to a loss of male-female courtship [12–14]. Fru^M^ is expressed in approximately 2000 interconnected neurons throughout the peripheral and central nervous system and its expression is required for the development, function, and plasticity of the circuit which drives male-specific behaviors [15]. In particular, social cues such as pheromones can affect courtship behaviors in males [16–21]. Two types of these pheromones, male-specific pheromone *cis*-vaccenyl acetate and non-sex-specific pheromones (such as methyl laurate and palmitoleic acid), activate Fru^M^-positive olfactory receptor neurons (ORNs) expressing Or67d and Or47b receptors, respectively [18, 20, 21]. These two ORN classes act differently, with Or67d regulating male-male repulsive behaviors and aggression, whereas Or47b driving age and social experience dependent male copulation advantage [4, 20–22]. Previous studies have reported that different social contexts, as well as loss of Or47b or Or67d function, alter the regulation of *fru* transcription, particularly the enrichment of active chromatin marks around *fru* promoters [23, 24]. In addition, the expression of *fru^M^* isoforms in Or47b and Or67d ORNs affects physiological responses to pheromone ligands and courtship behaviors [4, 20, 25, 26]. It is likely that changes in social context, pheromone signaling, as well as subsequent changes in *fru* regulation, affect the expression of ion channels as well as neurotransmitter receptors regulating neurophysiology. Indeed, Fru^M^ binding is detected upstream of many ion channels and genes controlling neural development and function in the central brain [27–29]. Even though these studies point to the regulation of neuronal and circuit function by Fru^M^, very little is known about how it affects the expression of these target genes, or how pheromone signaling and social experience affect transcriptional programs by modulating Fru^M^.

Here we performed antennal RNA-seq to determine transcriptional changes in response to social isolation and mutants in pheromone receptors or Fru^M^. Our results showed that *fru* splicing patterns were also modified in pheromone receptor mutants. We also found that transcriptional programs associated with neuromodulation were altered. Many of the Fru^M^ target neuromodulatory genes were misregulated in the same direction in *fru*^M^ and pheromone receptor mutants. These results uncover a gene regulatory cascade from pheromone receptors to *fru* regulation, which alters neuromodulatory transcriptional programs to ultimately modulate neuronal responses in different social contexts.

## Results

### Neuronal transcriptional programs are modulated with social isolation and lack of pheromone receptors or Fru^M^ function

To identify genes regulated in the peripheral olfactory system by social experience, pheromone signaling, and Fru^M^, we utilized RNA-seq from whole antennae of 7-day old wild-type (*w^1118^*) males that are either group-housed (*w^1118^ GH*) or single-housed (*w^1118^ SH*), as well as group-housed *Or47b* mutant males (*Or47b^1^*), *Or67d* mutant males (*Or67d^GAL4^*), and *fru^M^* mutant males (*fru^LexA^/fru^4-40^*) (Figure 1A). Each condition included 3 biological replicates except for *fru^LexA^/fru^4-40^* with only two (Figure 1B). Each sample had mapped reads ranging between 24 and 40 million and hierarchical clustering analysis based on Pearson’s correlation between samples showed consistency among replicates within the same genotype (Figure 1B). Principal component analysis (PCA) also showed the expected grouping of the replicates belonging to the same condition, across the first two principal components (PC) accounting for most of the overall variance (32% and 19%) (Figure 1C). We also found that gene expression changes were more similar among *Or67d, Or47b,* and *fru^M^* mutants, compared to grouped or isolated wild-type male antennae (Figure 1B,C). As expected, expression levels of *Or47b, Or67d*, and male-specific *fru* exon were significantly lower in all replicates for *Or47b*, *Or67d*, and *fru^M^*mutants, respectively, though the changes of the whole *fru* gene locus cannot be detected (Figure 1D, Figure 4B, Figure 4-figure supplement 1A), validating genotype-specific changes in each condition. In addition, genes known to be absent in adult antennae, such as *amos* [30–32], also showed nearly no expression, whereas housekeeping genes, like *Act5C*, *Gapdh2*, *RpII18*, *fl(2)d*, and *wkd*, showed nearly identical expression across all samples (Figure 1D). These results point to high RNA-seq data quality across sample groups and within biological replicates. The Or47b ORNs were shown to degenerate in 14-day old *Or47b* mutant flies. To test if the transcriptional changes are not due to the decrease of ORN numbers in 7-day old antennal samples, we counted the numbers of Or47b and Or67d ORNs. The total numbers of Or47b or Or67d ORNs were comparable between the control and the *Or* mutants (Figure 1-figure supplement 1). These results suggest that the changes of gene transcriptional level is mainly due to the loss of Or function, rather than the changes of ORN numbers.

**Figure 1.**
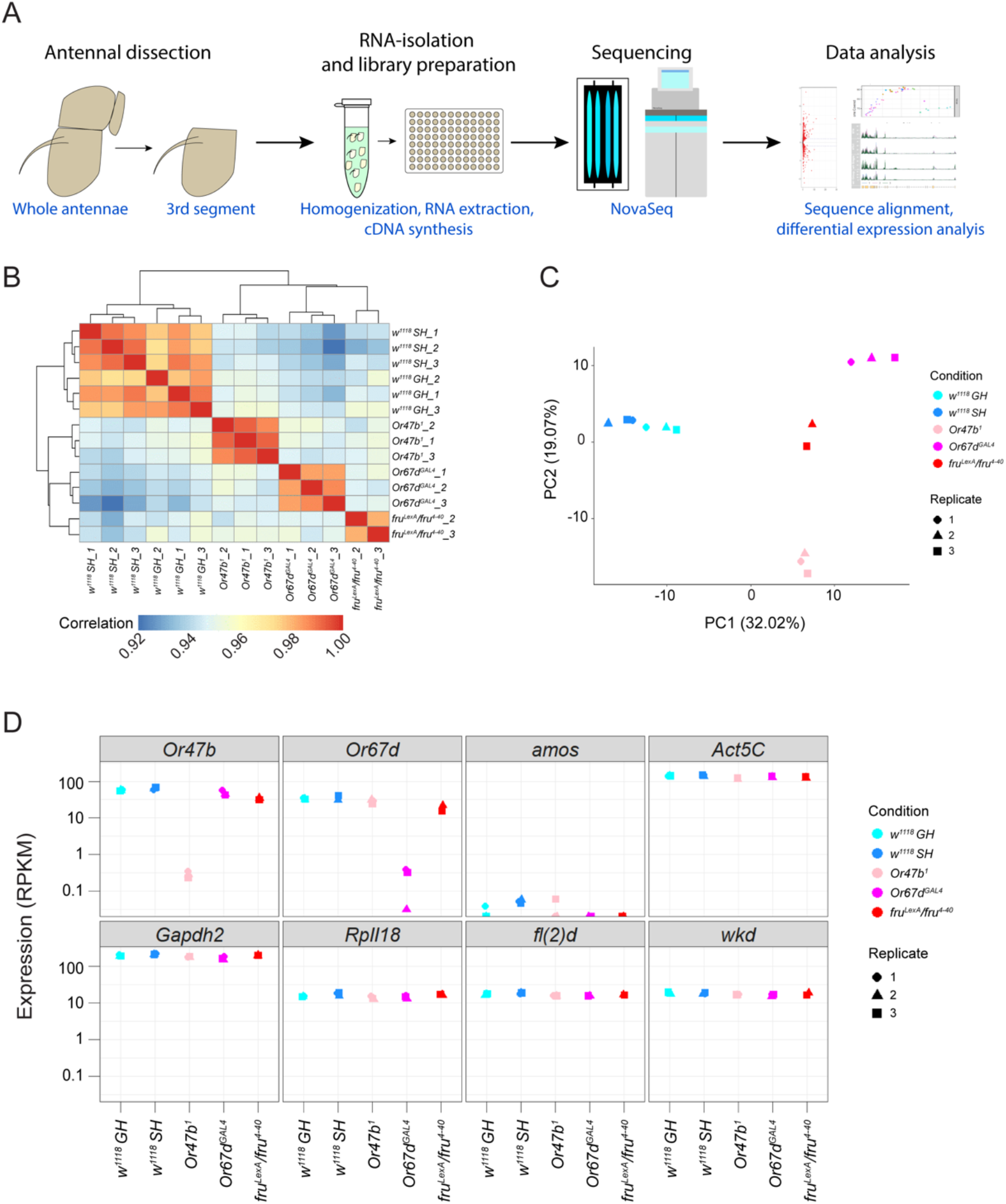
Overview of RNA-seq samples from male antennae. **(A)** schematic for antennal RNA-seq workflow. **(B-C)** Hierarchical clustering based on Pearson’s correlation matrix **(B)** and PCA analysis **(C)** of transcriptional profiles among biological replicates from antennae of wild-type group-housed (*w^1118^ GH*), single-housed (*w^1118^ SH*), and group-housed *Or47b^1^, Or67d^GAL4^,* and *fru^LexA^/fru^4-40^* mutant male flies. **(D)** Transcript levels for several representative negative and positive control genes among all samples.

**Figure 2.**
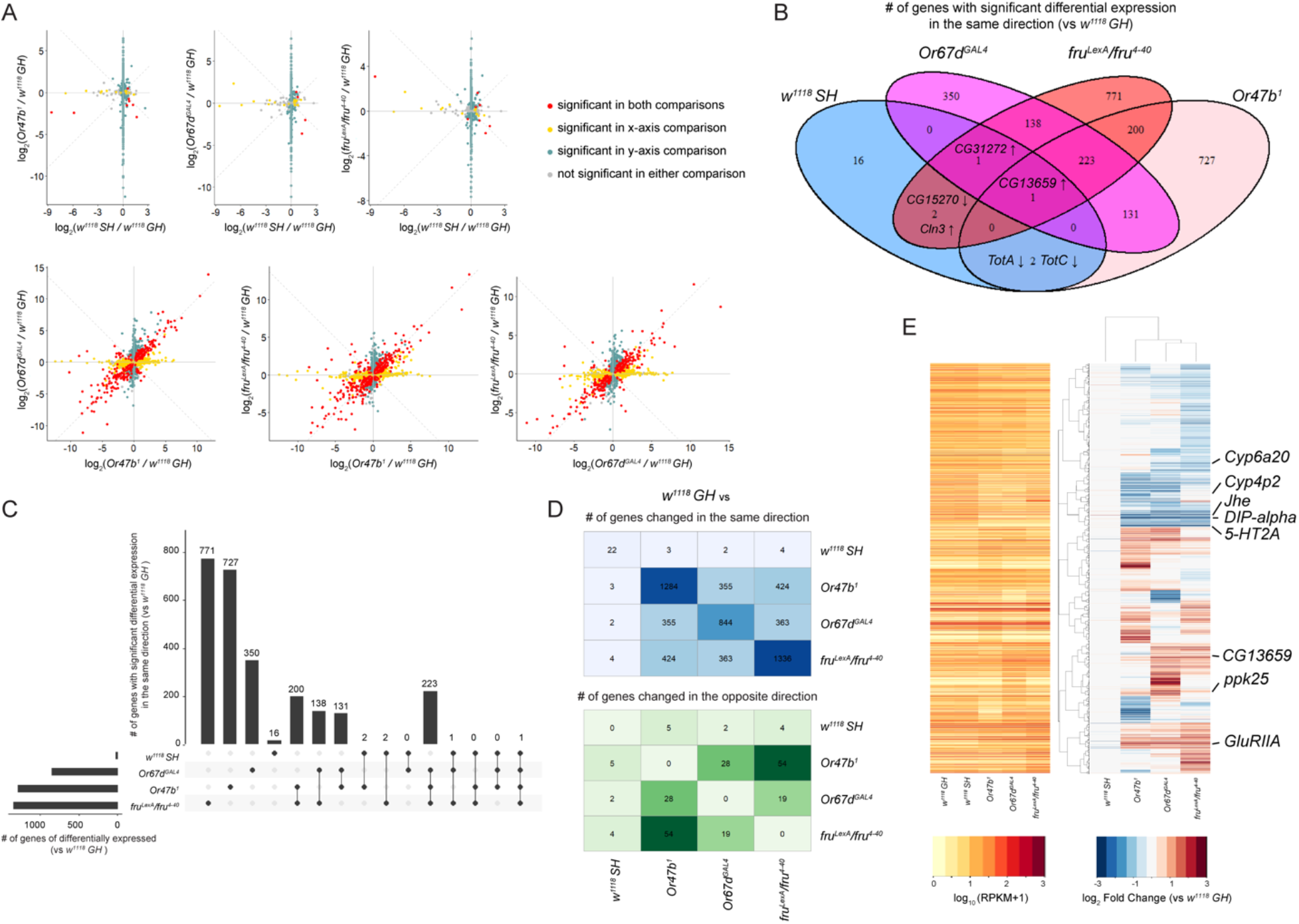
Differentially expressed genes in response to loss of social experience, pheromone receptors, or *fru^M^*. **(A)** Scatter plot showing the genes that are differentially regulated among social isolation, and mutants in pheromone receptors and *fru^M^*. Significance is defined by adjusted p-value below 0.01 after applying Bonferroni correction with n = 2. **(B-C)** Venn diagram **(B)** and UpSet plot **(C)** comparing differentially expressed genes shared across experimental conditions (only genes changed in the same direction). **(D)** Numbers of differentially expressed genes with the same (top) and the opposite (bottom) direction in pairwise comparison of experimental conditions versus group-housed wild-type samples. In (B-D), significance is defined by adjusted p-value below 0.01 after applying Bonferroni correction with n = 4. **(E)** Hierarchically clustered heatmaps showing log_2_ fold change compared to group-housed wild-type antennae across all experimental conditions (right) and average mRNA levels (reads per kilobase of transcript, per million mapped reads, RPKM) of replicates within each condition ordered in the same way as log_2_ fold change (left). Only 2999 genes with at least one significant (adjusted p-value below 0.01) change between an experimental condition versus group-housed wild types are shown.

**Figure 3.**
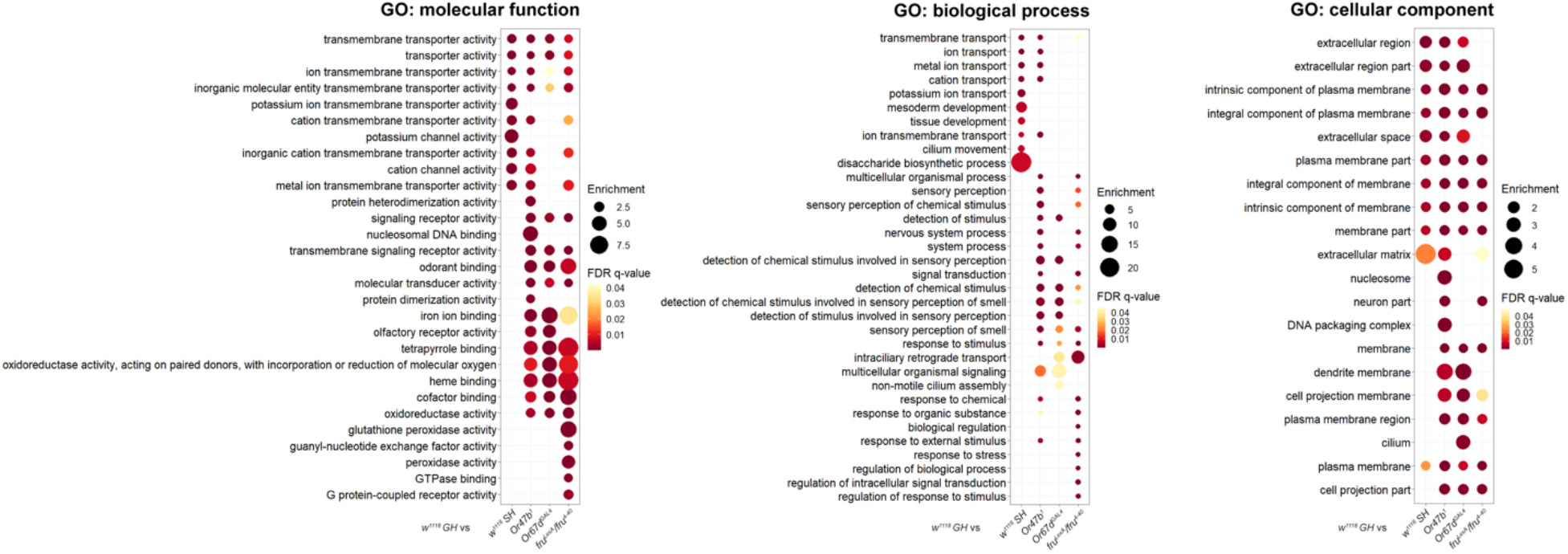
Top enriched gene ontology (GO) terms for differentially expressed genes in response to social experience, pheromone signaling, and Fru^M^ function. The union set of top 10 most significantly enriched GO terms with FDR q-value below 0.05 of the differentially expressed genes in each experimental condition is shown. Enriched GO terms were generated by the single ranked gene list with the most significantly changed genes at the top via GOrilla.

**Figure 4.**
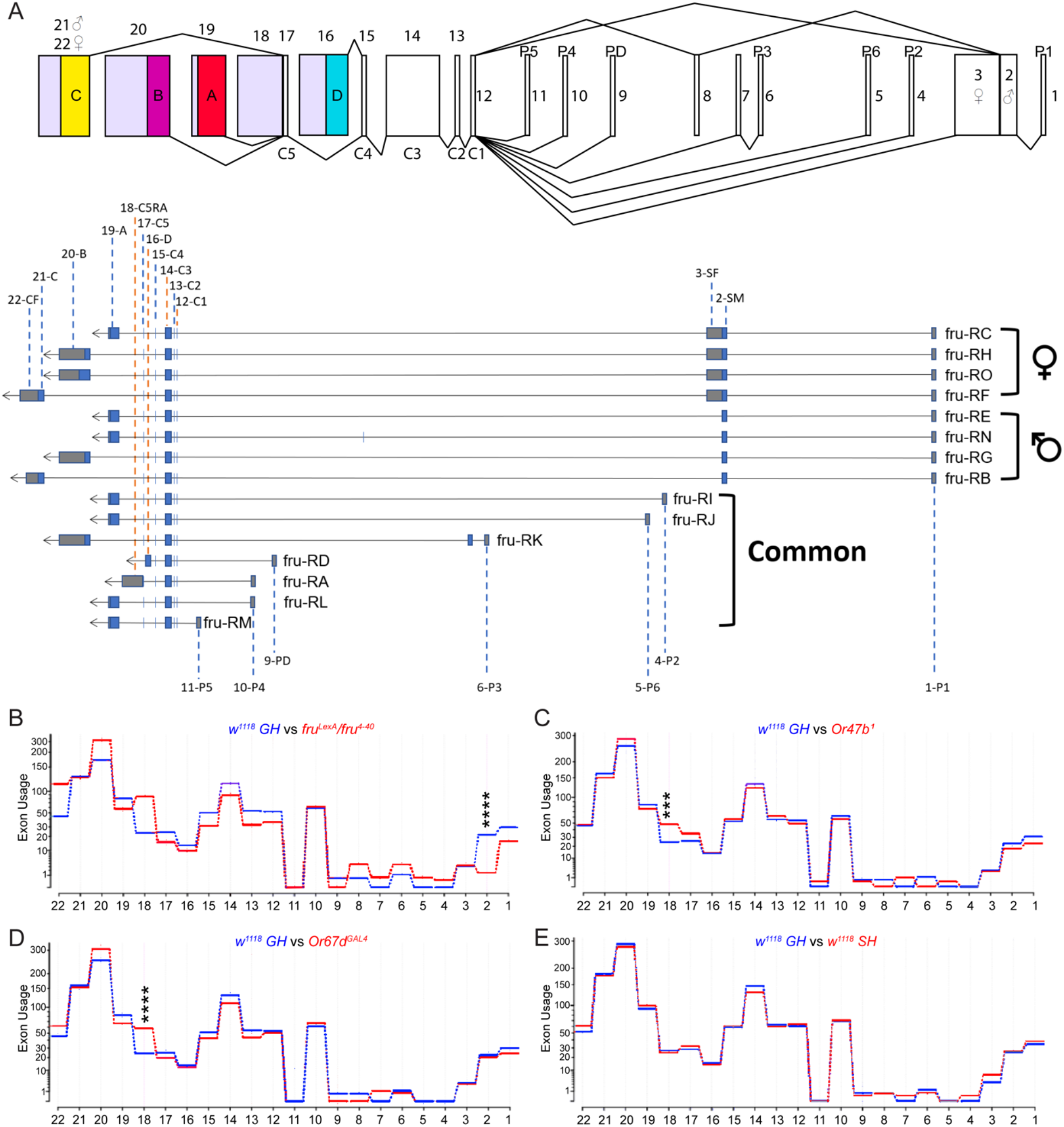
Olfactory stimuli regulate exon usage across the whole *fru* genomic locus. **(A)** Top: schematic of Fruitless: 1-22 denote each examined site. P1-P6 and PD denote promoters. C1-C5 denote common exons, while A-D denote alternatively spliced 3’ DNA binding-encoding domains. Bottom: structure of *fru* transcripts with the exon names and numbering used in the schematic above. **(B-E)** Examination of the usage of various exons in Fruitless to determine distinct changes in *fru^M^* transcripts using DEXSeq, showing exon-by-exon comparison of *w^1118^ GH* vs *fru^LexA^*/*fru^4-40^***(B)**, *Or47b^1^* **(C)**, *Or67d^GAL4^* **(D)**, and *w^1118^ SH* **(E)**. Adjusted p-value was directly performed via DEXSeq (see methods – statistical analysis). ***, p.adjust<0.001; ****, p.adjust<0.0001. Region 2, *w^1118^ GH* vs *fru^LexA^*/*fru^4-40^*, p.adjust=1.25×10^-10^; region 18 (3’UTR), *w^1118^ GH* vs *Or47b^1^*, p.adjust=8.17×10^-4^; region 18 (3’UTR), *w^1118^ GH* vs *Or67d^GAL4^*, p.adjust=2.00×10^-7^.

We then ran the differential expression analysis to globally examine the transcriptional changes upon loss of social expression, pheromone sensing, or Fru^M^ function. Compared to group-housed wild-type antennae, social isolation had the least number of significantly altered genes, whereas group-housed *fru^M^* mutants resulted in the highest number (Figure 2A-C, Supplementary table 1). Pairwise comparisons of group-housed wild types to isolated wild types, and *Or47b/Or67d/fru^M^*mutants revealed that the genes co-regulated by pheromone receptors and Fru^M^ tended to behave in the same direction in the corresponding mutants (Figure 2A,D), suggesting the shared downstream signaling pathways upon pheromone receptor activation and Fru^M^-dependent regulation. The numbers of genes with significant differential expression in the same direction shared by each condition compared to the group-housed wild types are illustrated in a Venn diagram and Upset plot (Figure 2B,C, Supplementary table 2), where genes with overlapping changes in social isolation and *Or47b*, *Or67d*, and *fru^M^* mutants are highlighted. Particularly, only one gene, *CG13659*, an ecdysteroid kinase-like domain encoding gene, is consistently changed across all experimental conditions compared to antennae from the group-housed wild-type males (Figure 2B).

Hierarchical cluster analysis of differentially expressed genes compared to group-housed wild-type samples showed that the transcriptional changes in *fru^M^* and *Or* mutants were most comparable with one another and most dramatically different from the control (Figure 2E). Single-housed wild types were most similar to group-housed wild types (Figure 2E). Cluster analysis identified several genes of behavioral, neuromodulatory, and developmental importance such as *Cytochrome p450 6a20* (*Cyp6a20*)*, serotonin receptor 2A* (*5-HT2A*)*, Juvenile hormone esterase* (*Jhe*), and *Dpr-interacting protein alpha* (*DIP-alpha*) (Figure 2E) [33–36]. Among these, antennal expression of *Cyp6a20,* which is downregulated in *Or47b*, *Or67d*, and *fru^M^* mutants, was previously shown to mediate effects of social experience on male-male aggression (Figure 2E) [33]. On the other hand, *Cyp4p2,* which is involved in hormone metabolism and insecticide detoxification [37–39], is only misregulated in *Or47b* mutants (Figure 2E). In addition to the downregulated genes, we also found some genes encoding ion channels and neurotransmitter receptors that were significantly upregulated (*ppk25* and *GluRIIa*) (Figure 2E). The heatmap for gene expression changes revealed gene clusters that were co-regulated by pheromone receptors and Fru^M^, in addition to gene clusters that were uniquely regulated by each OR and Fru^M^; this again highlights that the co-regulated genes tend to change in the same direction in pheromone receptor and *fru^M^*mutants.

### Gene ontology terms for differentially expressed genes in response to lack of social and pheromone signaling highlight neuromodulators

Previous work has demonstrated that social experience, pheromone signaling, and Fru^M^ activity can regulate the responsiveness of pheromone sensing ORNs to modify neuronal function and sex specific behaviors [4, 18, 20, 21, 33, 40]. To functionally understand system-level changes in gene expression with social isolation, lack of pheromone signaling, and *fru^M^* mutants, we next investigated gene ontology (GO) terms using GOrilla for the list of differentially expressed genes in each experimental condition in pairwise comparisons with group-housed wild types [41, 42] (Figure 3, Supplementary table 3). Many GO terms of molecular function and biological process were commonly affected across multiple experimental groups, suggesting the converging downstream molecular events in response to social experience and pheromone sensing mediated by Fru^M^ activity (Figure 3). Strikingly, the genes with the altered expression tended to be localized on the cell membrane (Figure 3, GO: cellular component) and have functions in ion transport across membrane (Figure 3, GO: molecular function), and appeared to be involved in the process of detecting and responding to olfactory stimuli (Figure 3, GO: biological process). This supports previous studies in providing a general mechanism for social experience, pheromone receptor signaling, and Fru^M^-dependent regulation of pheromone responsiveness of Or47b ORNs [4, 24, 25]. Furthermore, genes with oxidoreductase activity also had overlapping alterations across *Or47b*, *Or67d,* and *fru^M^* mutants, and many of these appeared to contribute to insect hormone metabolism (Figure 3, GO: molecular function). Interestingly, previous studies reported that juvenile hormone signaling works together with social experience in olfactory receptor neurons to modulate chromatin around *fru* locus [4, 24]. Our RNA-seq results also add an additional layer of complexity to hormone-social experience interactions, as social experience and pheromone signaling affects the levels of certain hormones by modifying hormone metabolism dynamics. In summary, social isolation, disrupted pheromone receptor signaling, and lack of Fru^M^ function in peripheral olfactory sensory neurons affect the expression of many genes with roles in diverse aspects of neurophysiology, including neuronal responsiveness, ion transmembrane transport, and beyond.

### Loss of pheromone signaling alters *fruitless* splicing patterns and *doublesex* expression

*Fruitless* locus generates multiple alternative splice isoforms for RNAs transcribed from 7 promoters (P1-P6, PD). The transcripts from *fru P1* are alternatively spliced between males and females, where the male, but not the female, isoform (*fru^M^*) encodes functional proteins (10,11). The expression of *fru^M^* in males and the absence of functional *fru^F^* transcripts in females help define male and female-specific neuronal pathways as well as the cell-specific expression patterns of genes regulated by Fru^M^. Promoter *fru P2* through *fru P6* produce common isoforms in both males and females that also affect sex-specific activity in courtship circuits of both sexes [43] (Figure 4A). *Fru^M^* itself has multiple splicing isoforms that vary in the 3’ end of the mRNA (*fru^MA^, fru^MB^, fru^MC^*), which encode proteins with variable zinc finger DNA binding domain of Fru^M^ transcription factor [27, 29, 43]. These regulate different aspects of the circuit controlling courtship behaviors, with Fru^MC^ and Fru^MB^ having the highest overlap behaviorally, and Fru^MA^ having little to no effect on courtship [27].

We previously showed that social experience and signaling from Or47b and Or67d pheromone receptors alter open chromatin marks around *fru P1* promoter in the male antennae [24]. Interestingly, examination of total transcript levels for the entire *fru* gene locus showed little to no difference across experimental conditions (Figure 1D). These small changes in total transcript levels, despite dramatic changes in open chromatin marks in wild-type SH, and mutants in *Or47b*, *Or67d,* and *fru^M^,* prompted us to look at other aspects of gene regulation. It is known that changes in chromatin regulate many aspects of transcription such as transcriptional initiation, elongation and alternative splicing [44, 45]. The effects of chromatin on splicing are thought to occur especially because chromatin state alters the speed of RNA Polymerase II (RNAPII), which can lead to splicing mistakes like intron retention or exon skipping [45].

Given the functional differences in the *fru^M^* isoforms, we predicted that chromatin changes caused by social experience and pheromone receptor signaling could alter *fru* splicing. To explore this, we mapped reads from all experimental conditions to *fru* genomic locus and investigated exon usage levels using DEXseq [46]. In general, transcript reads from *fru* locus appear noisier in experimental conditions compared to group-housed wild-type male antennae, with variations in the expression of coding and non-coding sequences (Figure 4B-E). In *Or47b* mutants, there is a small decrease in *fru P1* promoter (region 1, see methods – statistical analysis) and male-specific exon (region 2, see methods – statistical analysis) levels (Figure 4C). *Or67d* mutants show a small decrease in *fru P1* promoter levels (see methods – statistical analysis) and male-specific exon (region 2; see methods – statistical analysis) (Figure 4D). The largest change in male-specific exon levels is seen in *fru^LexA^/fru^4-40^* allele (Figure 4B), which has a *LexA* transgene insertion disrupting the male-specific exon and a 70-Kb deletion from the *P1* promoter [47]. *fru P1* promoter and the male-specific exon are unaltered in socially isolated antennae, yet there is a small increase in the female-specific exon (region 3, see methods – statistical analysis) (Figure 4E). In addition to the first 3 exons, a non-coding sequence (region 18) (Figure 4A), which is a only present in *fru-RA* transcript (43), strikingly increases in *Or67d, Or47b*, and *fru^M^* mutants (Figure 4B-D). This transcript encodes a Fru protein that lacks these zinc finger domains but retains BTB/PDZ protein-protein interaction domain (Figure 4A). It is possible that this isoform can interfere with the transcriptional functions of Fru^M^ proteins by binding and titrating out their interaction partners such as other transcription factors, chromatin modulators, and basal transcriptional machinery [48–51]. These results suggest social and pheromonal cues alter *fru* exon usage, likely indicating changes in *fru* splicing patterns. Another sex determination transcription factor known to regulate sex specific behaviors is *doublesex (dsx)* [52–60]. *dsx* expression in the antenna is restricted to non-neuronal cells [58]. We found that the expression of *dsx* in antenna is significantly increased in *Or* and *fru^M^* mutants, albeit the increase is much more pronounced in *Or67d* and *fru^M^* mutants (Figure 4-figure supplement 1B). Socially isolation did not alter the expression of *dsx* in antennae (Figure 4-figure supplement 1B,D). These results suggest that the expression of *dsx* in the antennae is repressed by *Or47b, Or67d* and *fru^M^*function.

In conclusion, our results suggest that the expression of two critical transcription factors, Fru and Dsx, that regulate sex-specific behaviors are modulated by pheromone signaling.

### Bimodal regulation of genes modulating neurophysiology and neurotransmission by Fru^M^ and pheromone receptor signaling

Previous studies have shown that pheromone receptor signaling and social experience-dependent regulation of chromatin and RNAPII enrichment around *fru P1* promoter can ultimately scale and fine-tune behavioral responses to social environment [4, 24]. Additionally, previous reports on the genome-wide binding profiles for three Fru^M^ isoforms in the central brain revealed isoform-specific differences in target genes that regulate neuronal development and function [27, 52]. Fru^M^ motifs are enriched among regulatory elements that are open in the female but closed in the male, suggesting Fru^M^ functions as possible repressive transcription factor [61]. Functional differences of Fru^M^ isoforms also influence ORN responses to their pheromone ligands [25]. Thus, chromatin-based modulation of *fru* levels and splicing with social experience and pheromone signaling can provide a quick way to modulate neuronal physiology and synaptic communication by modifying gene expression programs. Yet, the effect of social experience and pheromone receptor signaling on gene expression programs or the mode of gene regulation by Fru^M^ (as a transcriptional activator, repressor, or both) remains unknown. As discussed previously, gene ontology analysis of these differentially expressed genes implies that many genes involved in neuromodulation are regulated by social context, pheromone receptor signaling, and Fru^M^ function. To further investigate this, we specifically focused on genes associated with ion channel activity and/or neurotransmitter regulation (Figure 5A,B; Figure 6A,B). We clustered these genes based on their log_2_ fold change in transcript levels compared to group-housed wild types in each experimental condition, while also showing their corresponding expression levels in the antennae (RPKM, reads per kilobase of transcript, per million mapped reads) (Figure 5A,B; Figure 6A,B). We found many ion channel and/or neurotransmitter receptor-encoding genes showed up/down-regulation in response to social isolation, and loss of Or47b, Or67d, or Fru^M^ function (Figure 5A,B; Figure 6A,B). Within ion channels, two subclasses stood out. These are the Degenerin/Epithelial Sodium Channel (DEG/ENaC) proteins known as pickpockets (ppks) and inward-rectifying potassium channels Irks. Additional genes also include those encoding calcium channels, for example, *Piezo*, *TrpA1*, and *cacophony* (*cac)* (Figure 5A,B).

**Figure 5.**
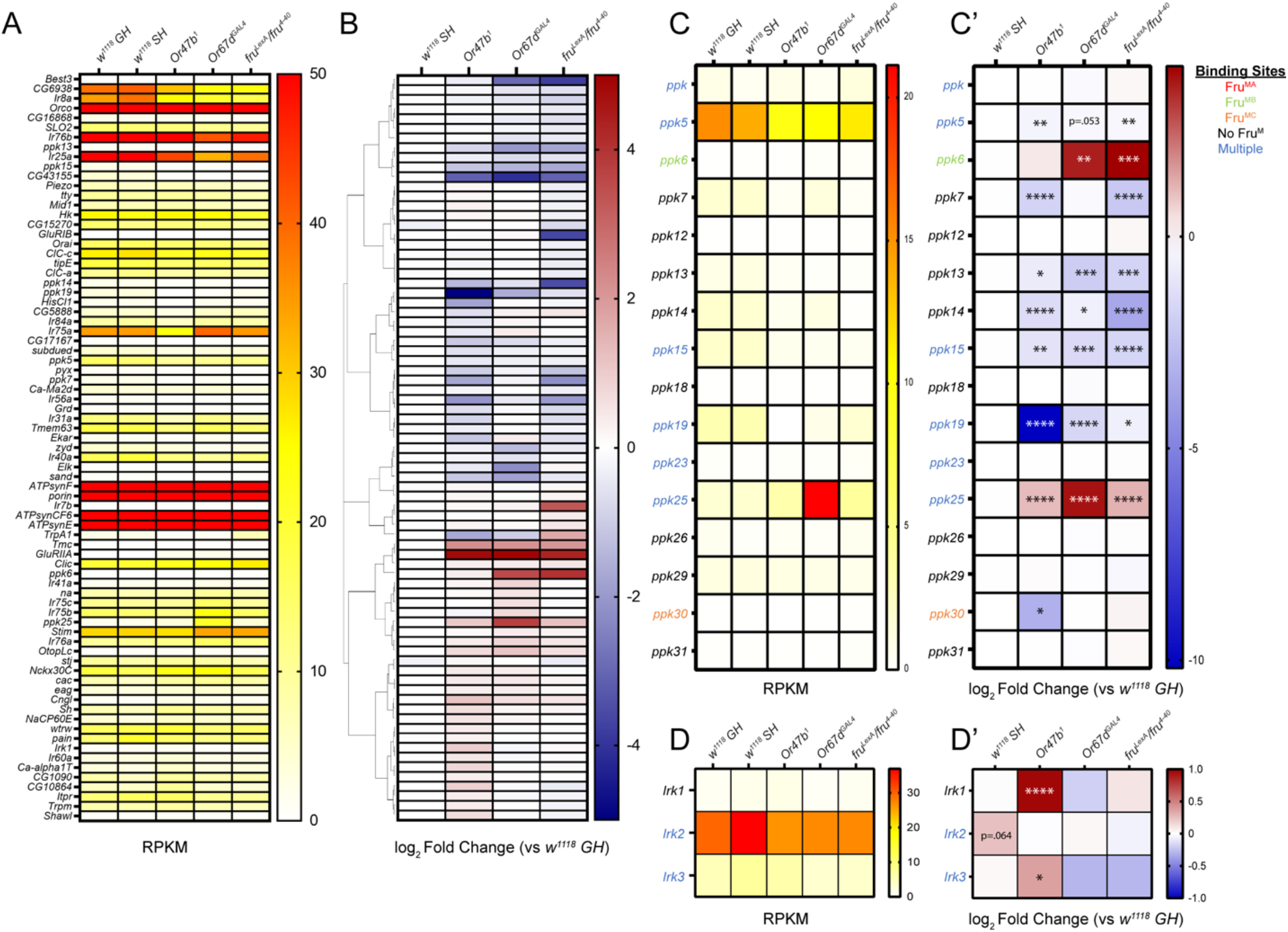
Differentially expressed ion channel-encoding genes in response to social isolation and loss of pheromone receptors or *fru*^M^. **(A-B)** Examination of GO term: 0005216 (ion channel activity) shows significant changes in various ion channel subclasses. Hierarchically clustered heatmaps showing log_2_ fold change compared to group-housed wild-type antennae across all experimental conditions **(B)** and average mRNA levels (RPKM) of replicates within each condition ordered in the same way as log_2_ fold change **(A)**. Genes with adjusted p-value above 0.01 were filtered out in each experimental condition. **(C-C’)** RPKM **(C)** and log_2_ fold change **(C’)** for *pickpocket* (*ppk*) gene family. **(D-D’)** RPKM **(D)** and log_2_ fold change **(D’)** for *inwardly rectifying potassium channel* (*Irk*) gene family. Adjusted p-value was directly performed via DESeq2. *, p.adjust<0.05; **, p.adjust<0.01; ***, p.adjust<0.001; ****, p.adjust<0.0001. Fru^M^ binding information is listed in Supplementary table 4.

**Figure 6.**
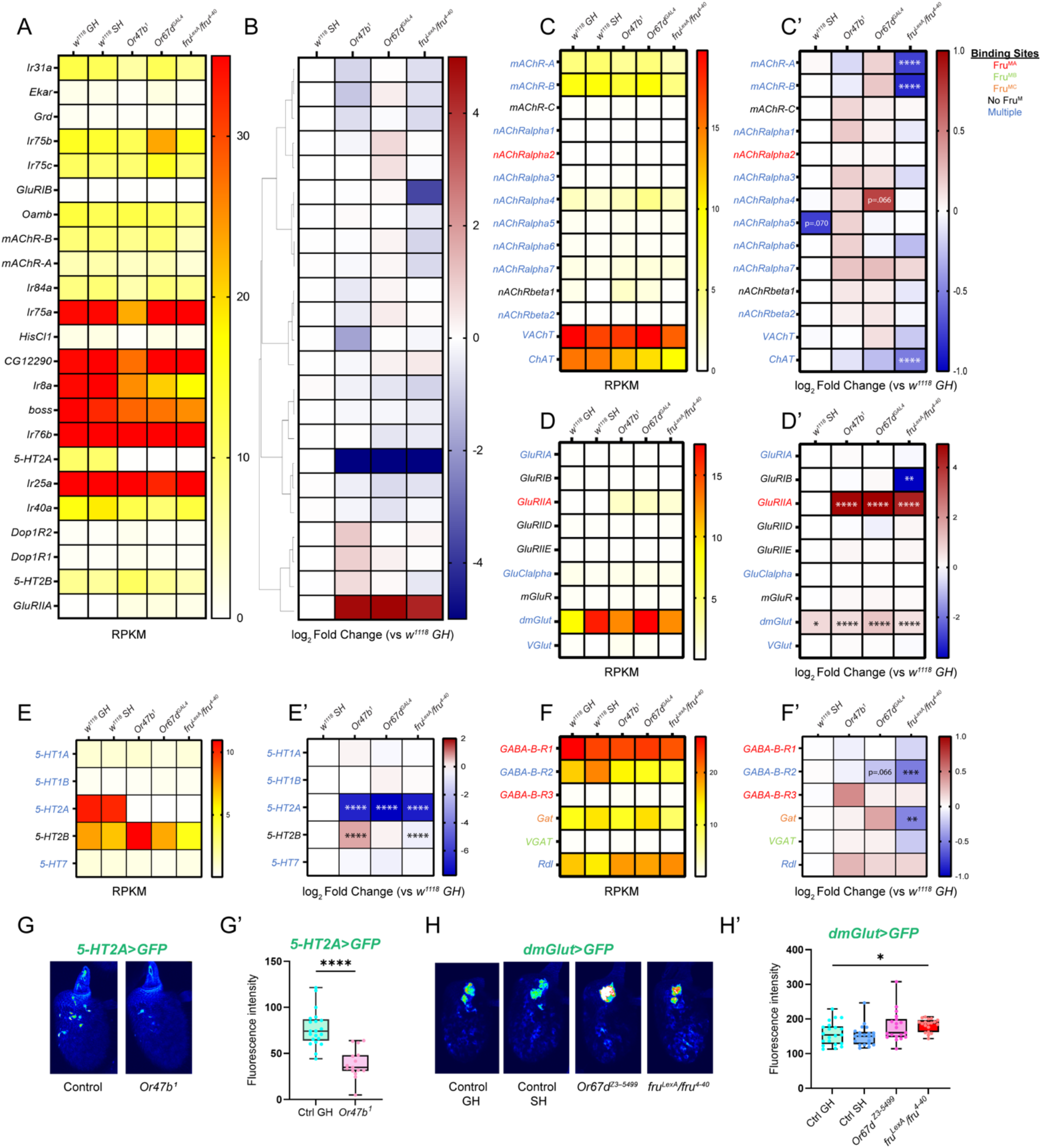
Differentially expressed neurotransmitter receptor and transporter-encoding genes in response to social isolation and loss of pheromone receptors or *fru^M^*. **(A-B)** Examination of GO term: 0030594 (neurotransmitter receptor activity) shows significant changes in various neurotransmitter activity-associated subclasses. Hierarchically clustered heatmaps showing log_2_ fold change compared to group-housed wild-type antennae across all experimental conditions **(B)** and average mRNA levels (RPKM) of replicates within each condition ordered in the same way as log_2_ fold change **(A)**. Genes with adjusted p-value above 0.01 were filtered out in each experimental condition. **(C-C’)** RPKM **(C)** and log_2_ fold change **(C’)** for acetylcholine-associated genes. **(D-D’)** RPKM **(D)** and log_2_ fold change **(D’)** for glutamate-associated genes. **(E-E’)** RPKM **(E)** and log_2_ fold change **(E’)** for serotonin-associated genes. **(F-F’)** RPKM **(F)** and log_2_ fold change **(F’)** for GABA-associated genes. Adjusted p-value was directly performed via DESeq2. *, p.adjust<0.05; **, p.adjust<0.01; ***, p.adjust<0.001; ****, p.adjust<0.0001. Fru^M^ binding information is listed in Supplementary table 4. **(G, H)** Confocal images of antenna from WT GH, WT SH, *Or47b^1^, Or67d^Z3-5499^,* and *fru^LexA/4-40^* mutants expressing *5HT2A-GAL4* driven *40XUAS-mCD8GFP* (**G)** and *dmGlut-GAL4* driven *UAS-mCD8GFP* (H). Quantification of fluorescence in images G **(G’)**, and H **(H’)**. **(G-G’)** n=15-21. **(H-H’)** n=19-21.

### ppk gene family

Recent reports pointed to the function of DEG/ENaC channels known as *pickpocket* family of sodium channels that act in Or47b and Or67d ORNs to regulate responses to their ligands [25]. Fru^M^ binding motifs have been identified around many of these *ppk* family members, such as *ppk*, *ppk5*, *ppk6, ppk15, ppk19*, *ppk23*, *ppk25,* and *ppk30* [27, 29, 62]. Both *ppk23* and *ppk25* have been identified as necessary for modulating responses of Or47b ORNs through Fru^MB^ and Fru^MC^ activity, respectively, with Fru^MB^ having an antagonistic effect on physiology in Or67d ORNs [25, 26]. We therefore first examined the expression levels of *ppk* genes in the antennae. In group-housed wild-type antennae, *ppks* show generally low expression, with *ppk5* displaying the highest levels (Figure 5C). Many *ppk* genes, including *ppk23,* seem not to be expressed in antenna. Analysis of recent single-cell RNA-seq data from ORNs [63] revealing ORN class-specific expression of *ppk*s also agree with this pattern. However, a few *ppks* known to be expressed in several ORN classes are not detectable in this dataset, possibly due to the limitations of single-cell RNA-seq in detecting low-abundance genes (Figure 5-figure supplement 1A). Even though there is no effect on the expression of any *ppk* family members in socially isolated male antennae, many *ppk* genes are differentially regulated in *fru^M^*mutants, in agreement with the existing Fru^M^ binding sites at their promoters. For example*, ppk6 and ppk25* are upregulated in *fru^M^*mutants whereas *ppk5*,*7*,*13*,*14*,*15*,*19* are downregulated. Many of these genes also show correlated changes in *Or47b* and/or *Or67d* mutants (*ppk13*,*14*,*15*,*19*,*25*). *ppk6* is strikingly upregulated in both *fru^M^* and *Or67d* mutants, whereas *ppk7* is downregulated in both *Or47b* and *fru^M^* mutants (Figure 5C’). Of note is the significant increase in *ppk25* expression, especially in *Or67d* mutants. *ppk25* is expressed in Or47b and Ir84 ORNs, but not Or67d ORNs, and has recently been shown to be downstream of Or47b and Ir84a activity altering their neuronal responses [26, 64, 65] (Figure 5C). The changes in *ppk25* levels were also validated through quantitative RT-PCR (Figure 5-figure supplement 1C-E). The bimodal changes in *ppk* transcripts in *fru^M^* mutants suggest Fru^M^ can act as both a repressor and an activator of *ppk* gene regulation. Furthermore, given that pheromone receptor function regulates Fru, the patterns of *ppk* gene misregulation in *Or47b* and *Or67d* mutants are likely dependent on changes in Fru^M^ activity downstream of pheromone receptor signaling.

### Irk gene family

Irk gene family encodes 3 inwardly rectifying potassium channels (Irk1-3) with binding motifs for Fru^MA^ identified upstream of *Irk2* and binding of both Fru^MA^ and Fru^MC^ found around *Irk3* [27, 29, 62]. Three *Irk* genes are expressed in varying levels in the antennae with *Irk1* having the lowest expression and *Irk2* having the highest expression (Figure 5D). We found that *Irk1* is upregulated in *Or47b* mutants, whereas *Irk2* shows the trend towards upregulation in response to social isolation (Figure 5D’).

These results suggest that changes in the transcript levels of Fru^M^-regulated sodium and potassium channels with social isolation and in pheromone receptor mutants may contribute to changes in neuronal responses and behaviors.

#### Regulators of Neurotransmission

To ask if social experience, pheromone signaling, and Fru^M^ function regulate genes involved in neurotransmission, we next examined the expression of neurotransmitter receptors, transporters, and enzymes for neurotransmitter metabolism. ORNs in the antennae as well as their projection neuron targets around the antennal lobes are mostly cholinergic [67]. In the antennal lobe it has been shown that local interneurons, which include serotonergic, GABAergic, and glutamatergic interneurons, provide cross talk between synaptic partners in the antennal lobe glomeruli [67, 68]. These neurons form connections with both presynaptic ORNS and their postsynaptic partner projection neurons for modulation of neuronal response across glomeruli [67, 69, 70]. These connections are required for fine tuning of signaling at synapses as a way of rapid modulation of neuronal function [34, 69–77]. We found a high expression of choline acetyltransferase (*ChAT*) that catalyzes acetylcholine biosynthesis and *VAChT* that packages acetylcholine into synaptic vesicles, coinciding with their reported cholinergic roles in ORNs. Moreover, we also found relatively high expression of several genes encoding receptors of various neurotransmitters, such as choline, serotonin (5-HT), GABA, and glutamate (Figure 6C-F’). Many of these genes, such as *nAChRalpha4*/*5*, *5-HT2A*, *5-HT7*, *GABA-B-R2*, and *GluRIIA*, have previously been found to regulate courtship behavior in flies through signaling in the antennal lobe [73, 78–80]. Interestingly, *GABA-B-R2* was shown to be specifically involved in presynaptic gain control of Or47b ORNs [81]. Additionally, single-cell RNA-seq data shows both broadly expressed neurotransmitter genes like *GluRIIB* and 5-*HT2B*, while others are specific to a subset of ORN classes [66] (Figure 6-figure supplement 1). Overall, many of the genes encoding neurotransmitter receptors show expression changes in different experimental conditions (Figure 6B).

To focus on genes related to specific neurotransmitters, we didn’t observe any significant changes in response to social isolation, except for a few genes, like *dmGlut,* which is upregulated compared to the group-housed wild types (Figure 6D,D’). We again found that loss of Fru^M^ function led to bimodal effects on gene expression (Figure 6C-F’). Indeed, many of these genes have known Fru^M^ binding to their promoters, including receptors *nAChRalpha1*/*3*/*4*/*5,* G*luRIIA*, *GluClalpha*, *5-HT1A*, *5-HT1B*, *5-HT2A*, *5-HT7*, and transporters/regulators such as *VAChT, ChAT*, and *Gat* [27, 29, 62]. Some of these genes display correlated changes between pheromone receptor mutants and *fru^M^* mutants, like *GluRIIA*, *dmGlut*, and *5-HT2A*, suggesting that the effects of pheromone signaling on neurotransmission can act via their influences on *fru* regulation (Figure 6D-E’). The changes in *5-HT2A* were also validated through qRT-PCR (Figure 6-figure supplement 1C). In the antenna, *5-HT2A*-*GAL4* and *dmGlut-GAL4* expression is observed in a subset of ORNs (Figure 6G-H’). Interestingly, Or47b and Or67d ORNs do not express *5-HT2A* reporter (Figure 6G). In agreement with a decrease in *5-HT2A* transcript levels in the RNA-seq and RT-PCR experiments, *5-HT2A* reporter expression was significantly decreased in *Or47b* mutant antennae (Figure 6G-G’). On the other hand, *dmGlut* expression in the antennae was upregulated in all conditions compared to group housed male antennae, generally in agreement with the qRT-PCR validation results (Figure 6D,D’, Figure 6-figure supplement 1D). We also used a *dmGlut-GAL4* to visualize the *dmGlut* expression in vivo, and detected the signal in a subset of non-neuronal cells in the antennae, though we only observed the statistically significant increase of the *dmGlut* reporter expression in *fru^M^* mutants (Figure 6H,H’). Evident changes are also observed in some genes not known to be Fru^M^ targets, for example, *GluRIB* which shows downregulation only in *fru^M^*mutants, and *5-HT2B* which shows upregulation in *Or47b* and *fru^M^*mutants (Figure 6D-E’). These may reflect effects of pheromone receptor signaling independent of Fru^M^ function or indirect effects of Fru^M^ activity. To summarize, the systems-level changes in expression of genes involved in neurotransmission and neurophysiology with social experience and pheromone receptor signaling can modulate ORN responses. In addition, these effects on gene expression with social signals can occur either in a Fru^M^-dependent manner or independently of Fru^M^ in response to other gene regulatory pathways activated by pheromone receptor signaling.

#### Odorant binding proteins

Analysis of GO terms for molecular function for “odorant binding” highlighted genes encoding odorant binding proteins (Obps) among the significantly altered compared to group housed wild type male antennae (Figure 7). Previous studies using in situ hybridization and transcriptional reporters have shown that Obps are generally produced in the non-neuronal support cells of the antennal sensilla and are secreted into the local hemolymph [82]. However, our analysis of previously published single-cell RNAseq data from ORNs [66] revealed that some Obps (i.e. *Obp19a, Obp28a, Obp56a, Obp59a, Obp69a, Obp83a, Obp83b,* and *lush)* are abundantly expressed in ORNs as well (Figure 7-figure supplement 1A). Odorants that enter the sensilla through pores are thought to interact with Obps in the hemolymph, which aid with odor binding to receptors on the cilia of ORNs [82]. Mutants in Obp genes are associated with alterations in the spontaneous or evoked neuronal response dynamics of resident ORNs [82–87].

**Figure 7.**
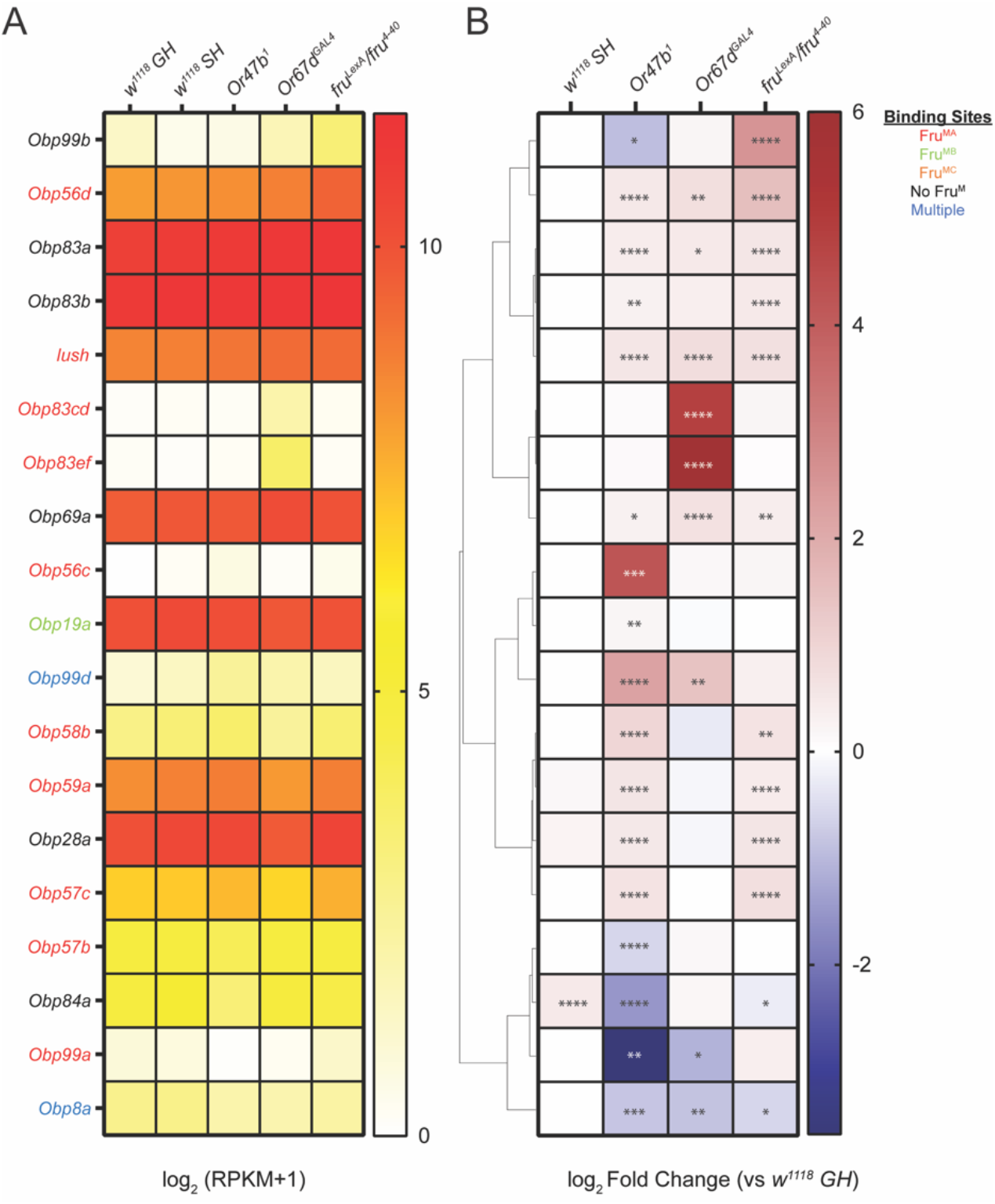
Differentially expressed *Odorant binding protein (Obp)* genes in response to social isolation, loss of pheromone receptors or *fru^M^*. **(A-B)** Many *Obp* genes show significant changes. Hierarchically clustered heatmaps showing log_2_ fold change compared to group-housed wild-type antennae across all experimental conditions **(B)** and average mRNA levels (RPKM) of replicates within each condition ordered in the same way as log_2_ fold change **(A)**. Genes with adjusted p-value above 0.01 were filtered out in each experimental condition. Adjusted p-value was directly performed via DESeq2. *, p.adjust<0.05; **, p.adjust<0.01; ***, p.adjust<0.001; ****, p.adjust<0.0001. Fru^M^ binding information is listed in the supplementary file. Fru^M^ binding information is listed in Supplementary table 4.

Analysis of Obp gene expression in the mutant male antennae showed many Obp transcripts that normally are expressed in trichoid sensilla were increased in the antennae from *Or47b, Or67d* and *fru^M^* mutants (eg. *Obp83a, Obp83b, lush, Obp69a*) [82, 85–89] (Figure 7). qRT-PCR from antenna generally corroborates RNA-seq results (Figure 7-figure supplement 1B,C).

Among the Obps that are differentially expressed in mutants, *Obp69a* is particularly interesting as it was previously shown to modulate social responsiveness in Drosophila [90]. In this context, cVA exposure in males as well as activation of Or67d neurons decreases *Obp69a* levels, which in turn alters aggressive behaviors driven by Or67d neurons. In addition, *lush*, *Obp83a* and *Obp83b* which are also expressed in trichoid sensilla were all shown to regulate odor evoked response kinetics and spontaneous activity of trichoid ORNs [83, 85–87].

In addition to Obps expressed in the trichoid sensilla, many other Obps also show misregulation particularly in *Or67d* and *Or47b* mutants. For example, in both mutants *Obp99d* is significantly upregulated; in contrast, *Obp99a* and *Obp8a* show a downregulation (Figure 7). Even though it is not known which sensilla these Obps are normally expressed in, given the responses it is likely that they are produced by the non-neuronal cells in trichoid sensilla where Or47b and Or67d ORNs are housed. There are also some Obps that show misregulation only in specific mutants. For example, *Obp83cd, Obp83ef, Obp56c* are normally not expressed in the antennae, yet *Obp83cd, Obp83ef* show a significant upregulation in *Or67d* mutants, whereas *Obp56c* is upregulated in *Or47b* mutants (Figure 7). *Obp84a* is the only *Obp* to be upregulated in isolated male antennae, and downregulated in *Or47b* mutant antennae (Figure 7). These results suggest the presence of regulatory interactions between olfactory receptor signaling and neural activity that likely drive activity-dependent homeostasis in Obp levels. Given the role of most Obps in regulating neuronal physiology, it is possible that transcriptional changes in *Obp* genes observed in social isolation as well as pheromone receptor mutants might occur as homeostatic mechanism to compensate for altered neuronal activity and ORN function.

### Pheromone receptor signaling regulates genes involved in hormone metabolism

Hormone signaling is responsible for regulating behavioral and brain states in both vertebrates and invertebrates. For example, in vertebrates many social behaviors such as aggression, mating, and parenting, are under the control of hormones such as estrogen, testosterone, oxytocin and vasopressin [91–94]. In social insects, such as ants, caste-specific behaviors are determined by hormone states, where queen- and worker-like behaviors are associated with ecdysone and juvenile hormone signaling, respectively [95, 96]. In Drosophila, juvenile hormone signaling modulates behavioral and motivational states during courtship [20, 97, 98]. Recent studies have also identified age-related cues such as juvenile hormone (JH) signaling together with social experience to control Or47b neuronal responses to pheromones and courtship behaviors in a Fru^M^-dependent manner [4, 20, 25]. JH signaling, concurrent with social experience, modifies chromatin around *fru P1* promoter and ultimately *fru^M^* levels in Or47b ORNs [24]. These studies also demonstrated that JH receptor enrichment at *fru P1* promoter increases in socially isolated flies as well as flies with disrupted Or47b signaling [24]. As mentioned above, gene ontology analysis of differentially expressed genes in this study also highlights genes involved in hormone metabolism (Figure 3). Thus, we specifically interrogated the genes regulating hormone levels in pheromone receptor and *fru^M^*mutants (Figure 8A-C’). Many of the enzymes involved in juvenile hormone biosynthesis and metabolism, such as juvenile hormone epoxide hydrolases (*Jheh1*,*2*,*3*), juvenile hormone esterase (*Jhe*), and juvenile hormone acid methyltransferase (*jhamt*), are expressed at varying levels in the antennae (Figure 8C). These genes are also reported to have Fru^MA^ and Fru^MC^ binding in their upstream regulatory elements (Dalton et al., 2013; Neville et al., 2014; Vernes, 2014). Two mostly enriched genes, *Jheh1* and *Jheh2*, show mild upregulation in *fru^M^* mutants but no significant changes in the absence of social cues or pheromone receptor signaling (Figure 8C,C’). On the other hand, both *Jhe* and *Jheh3* appear to be upregulated in social isolation while downregulated in *Or47b* mutants (Figure 8C’; Figure 8-figure supplement 1B).

**Figure 8.**
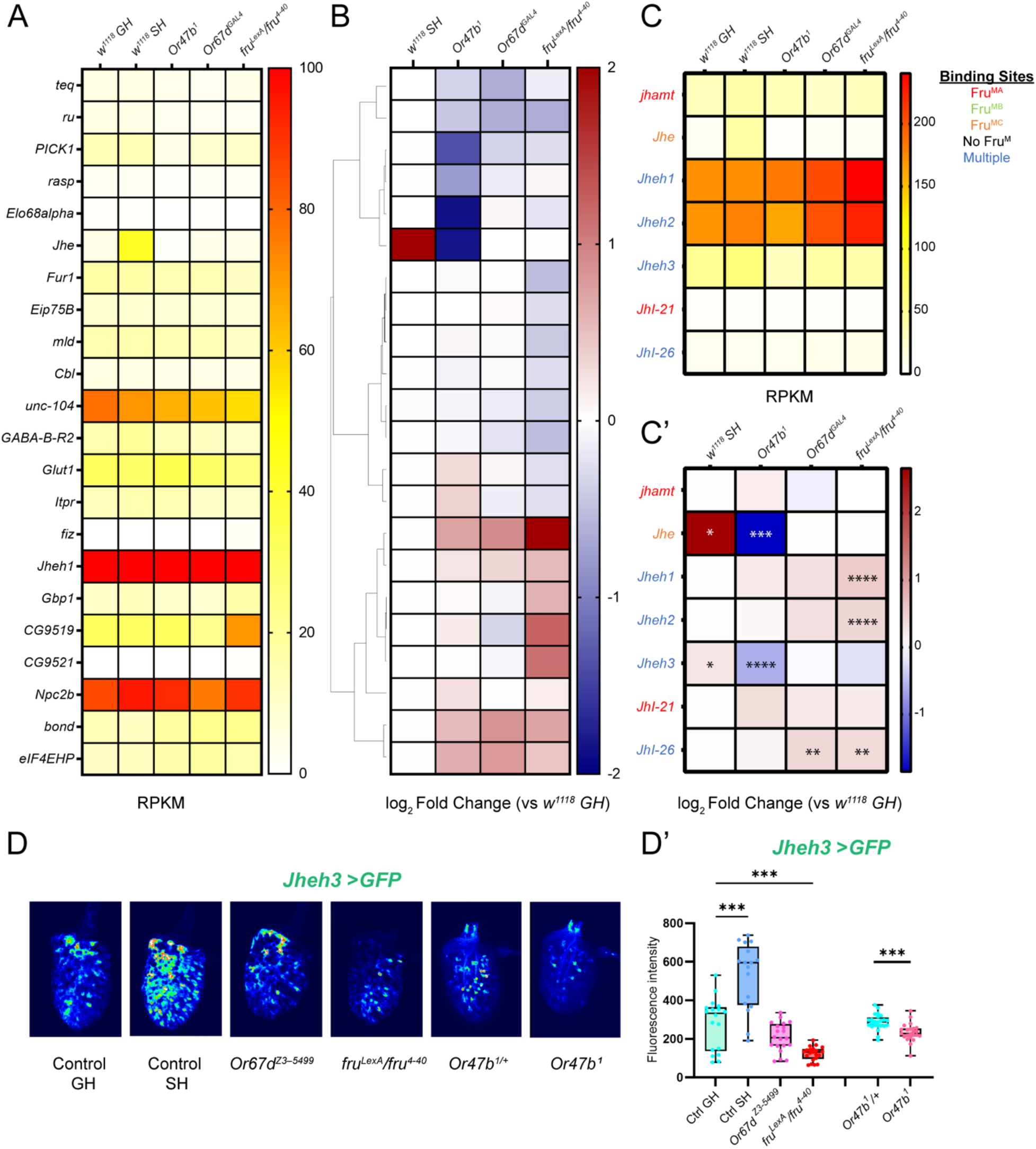
Differentially expressed hormone metabolism genes in response to social isolation and loss of pheromone receptor or *fru^M^*. **(A-B)** Examination of GO term: 0010817 (regulation of hormone levels) shows significant changes in various hormone metabolism gene subclasses. Hierarchically clustered heatmaps showing log_2_ fold change compared to group-housed wild-type antennae across all experimental conditions **(B)** and average mRNA levels (RPKM) of replicates within each condition ordered in the same way as log_2_ fold change **(A)**. Genes with adjusted p-value above 0.01 were filtered out in each experimental condition. **(C-C’)** RPKM **(C)** and log_2_ fold change **(C’)** for juvenile hormone metabolism-related genes. Adjusted p-value was directly performed via DESeq2. *, p.adjust<0.05; **, p.adjust<0.01; ***, p.adjust<0.001; ****, p.adjust<0.0001. Fru^M^ binding information is listed in Supplementary table 4. **(D)** Confocal images of antenna from WT GH, WT SH, *Or47b^1^, Or67d^Z3-5499^,* and *fru^LexA/4-40^* mutant males expressing *Jheh3-GAL4 UAS-mCD8GFP.* (D’) Quantification of fluorescence in images presented in D. **(D-D’)** n=17-24.

Throughout the antenna, *Jheh3*-*GAL4* expression is observed in many ORNs (Figure 8D, D’). In agreement with a transcriptional increase in socially isolated male antennae we found that *Jheh3* reporter expression was significantly increased in isolated male antennae. On the other hand, in both *Or47b* and *fru^M^*mutant antennae *Jheh3* reporter expression was significantly decreased in agreement with transcript levels (Figure 8D,D’). As observed in the RNA-seq, there was no change in *Jheh3* reporter expression in *Or67d* mutants compared to grouped male antennae (Figure 8D,D’). *Jhe* is of particular interest as Jhe activity is known to be necessary for robust male-specific courtship behaviors and mating success in addition to affecting the abundance of sex-specific pheromones such as 11-cis-vaccenyl acetate in males [36, 99].

Furthermore, seminal work on Jhe and Jheh3 has shown that these enzymes work together to catabolize JH in *D. melanogaster* [100]. These results suggest that social experience and pheromone receptor signaling regulates the expression of JH biosynthetic enzymes. Such changes can modulate juvenile hormone activity by rapidly catabolizing JH in the periphery and affecting downstream target genes, such as *fruitless*.

## Discussion

Sensory experience influences many behaviors by modifying neuronal and circuit function [1–4], yet, molecular mechanisms remain largely unknown. Here, we took advantage of the well-characterized system of sex-specific behaviors, governed by the Fru^M^, which acts as a gene regulatory switch for male-specific circuit development, function, and behavior in *Drosophila melanogaster* [10, 11, 15, 101]. Our results show that social experience and pheromone signaling alters Fru^M^ splicing patterns and neuromodulatory gene expression programs, which ultimately modulate circuit function and behavioral responses (Figure 9).

**Figure 9.**
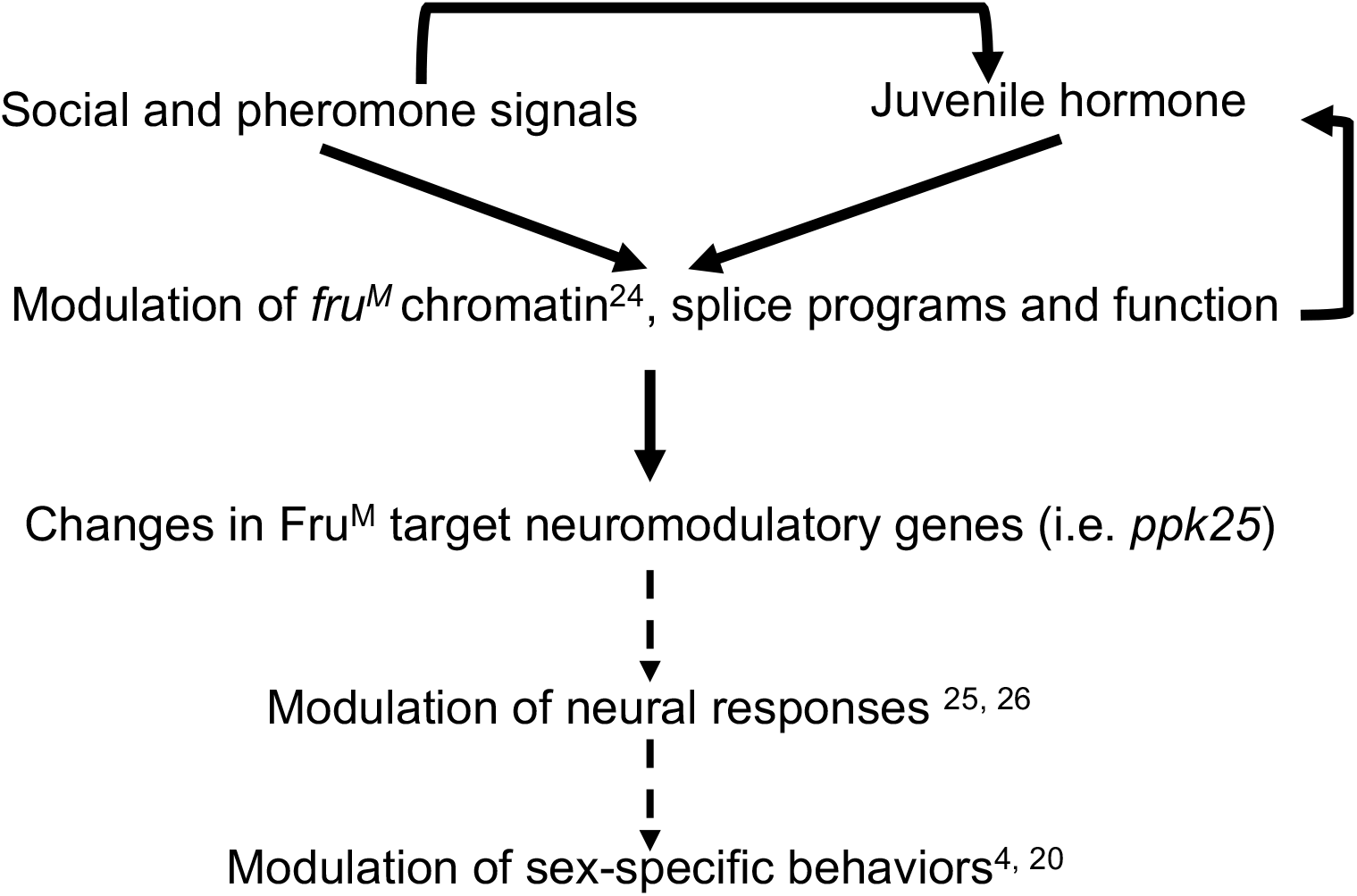
Fruitless-dependent transcriptional cascade that reprograms neural responses and behaviors with social experience, pheromone receptor function and hormone signaling. Social context and pheromone detection modifies chromatin [24] and transcriptional/splice programs for *fruitless* gene altering its function. This reprograms expression of Fruitless target neuromodulatory genes (i.e. *ppk25*) altering neural physiology and pheromone responses [4, 25, 26]. Ultimately, these result in changes in neuronal activity and behavioral modulation [4]. It was also shown that juvenile hormone signaling works together with social experience to modulate both ORN physiology and courtship behaviors [20, 25]. At the molecular level social/pheromonal cues work together with juvenile hormone receptors to modulate transcription *fruitless* [24]. Social context, pheromone receptor, and Fru^M^ function also alter the expression of genes involved in juvenile hormone metabolism.

Previous studies in Drosophila demonstrated that social experience can modulate Fru^M^-dependent sex-specific behaviors such as courtship and aggression [1, 3, 4]. For example, social isolation decreases the sensitivity of Or47b neurons to their pheromone ligands in a Fru^M^-dependent manner, which leads to a decrease in male competitive courtship advantage [4]. Other studies have also shown that monosexual group housing can decrease aspects of courtship behaviors such as courtship song and circling [102]. In addition to courtship, aggression behaviors which are under the control of Or67d and Or65a neurons and Fru^M^ function, also change with social experience [40, 102]. For example, social isolation significantly increases male-male aggression [33, 102]. These reports highlight the importance of social experience and pheromone signaling in the execution of sex-specific behaviors.

What are the molecular mechanisms by which Fru^M^ function is altered by social experience? We previously reported that social experience and pheromone receptor signaling alter chromatin states around *fru* P1 promoter [24] to modify *fru* transcription [4, 23, 24]. Nonetheless, the consequential details of chromatin-based changes on *fru* transcription remained unknown. One of our most intriguing findings from this transcriptome analysis is that these chromatin effects are associated with changes in the relative exon usage and splicing patterns of *fru* gene in response to impaired pheromone detection (Figure 4). Transcriptional regulation of *fru* is complex yielding 15 annotated alternatively spliced isoforms from 7 promoters giving rise to different 3’ sequences which encode variable Zinc-finger DNA-binding domains of Fru protein [13, 27, 103, 104]. Different Fru proteins regulate unique yet overlapping set of target genes which have binding sites for single or multiple Fru^M^ isoforms [27, 29, 62]. Many of these target genes regulate neural development and function. Therefore, changes in *fru* splicing patterns can affect the expression of thousands of genes simultaneously, strongly modulating neuronal responses and circuit outputs in a short period of time. Remarkably, Fru^M^ is expressed in ∼2000 interconnected neurons highlighting a circuit for courtship behaviors from sensation to action [11, 105]. This expression pattern allows neural activity-dependent influences on *fru* chromatin and transcription to propagate throughout the whole circuit. In summary, these features make circuit switch gene *fru^M^* an efficient molecular hub onto which many internal and external states act to modulate circuit activity and behavioral outputs by tweaking the levels of transcripts and splice isoforms, leading to a cascade of changes in transcriptional programs.

Each pheromone sensing neuron relays different information about the social environment, which is integrated and processed to output a specific behavior. Likely due to differences in neuronal identity and function, different pheromone receptors have different effects on *fru* chromatin and splice isoforms [24] (Figure 4). Such sensory stimuli-dependent changes in Fru proteins can alter the expression of downstream neuromodulatory genes to have rapid, temporary, or lasting effects on neuronal activity and behavioral outputs. These changes are essential for organisms to form short/long-term adaptation to the environment. However, how these different cell types generate these differences in behavioral repertoire via changes in gene expression in the periphery have been largely unknown.

Many of the genes that show differential expression in response to social isolation and disruption of pheromone receptor or Fru^M^ function encode neuromodulators that affect membrane potential, such as ion channels, membrane ion transporters, proteins involved in neurotransmission, and odorant binding proteins (Figure 3; Figure 5; Figure 6). Among all conditions, social isolation possesses the fewest differentially expressed genes compared to group-housed controls with a small overlap with pheromone receptor and *fru^M^*mutants. This might be due to differences in gene expression changes in response to disruption of evoked activity of pheromone sensing olfactory neurons with socially isolation versus disruption of both spontaneous and evoked activity in pheromone receptor mutants. Loss of Fru^M^ alters the expression of many neuromodulatory genes with known Fru^M^ binding sites in a bimodal way, suggesting Fru^M^ can act as both an activator and repressor of gene expression. Some of these differentially expressed genes are also altered in pheromone receptor mutants, generally in the same direction (Figure 2D,E). There are also unique overlaps between *Or47b* and *fru^M^* mutants, between *Or67d* and *fru^M^* mutants, and between *Or47b* and *Or67d* mutants (Figure 2B,E). Many of these differentially expressed genes are known to harbor binding sites for different Fru^M^ isoforms. These suggest that some of the differentially expressed genes in *Or47b* and *Or67d* mutant are due to Fru^M^-dependent changes, whereas others might be Fru^M^-independent, caused by OR signaling and/or ORN activity.

One functionally relevant gene among the genes that show differential regulation in pheromone receptor and *fru^M^* mutants is the Fru^M^ target gene *ppk25*, which previously was shown to modulate ORN responses in Or47b and Or67d neurons [25, 26]. *ppk25* belongs to a family of sodium channels that serve a variety of functions, from neuromodulation to detection of sensory cues. PPK protein complexes, generally are composed of multiple subunits encoded by different *ppk* genes. Many *ppk* genes contain binding sites for Fru^M^ isoforms in their promoter regions [27, 29, 62]. In addition, a recent study implicated isoform-specific Fru^M^-dependent regulation of *ppk25* and *ppk23* in the modulation of Or47b and Or67d responses [25, 26]. According to the genetic analysis in this study, Fru^MB^ and Fru^MC^ positively regulate the expression of *ppk25* and *ppk23*, respectively. There are apparent discrepancies with this interpretation and transcriptome data from our study, as well as others [66, 106]. While our transcriptome analysis agrees with a regulatory role for Fru^M^ in *ppk25* gene regulation, the regulatory mode is repressive; that is, *ppk25* expression is upregulated in *Or47b*, *Or67d,* and *fru* mutants. This type of repressive role for Fru^M^ in transcription also is in consensus with previous studies demonstrating Fru^M^ interactions with transcriptionally repressive histone-modifying enzymes such as HDAC1 [48, 107]. In addition, we are not able to detect any transcripts for *ppk23* in the antennae, and the expression of *ppk23* does not change in *Or47b*, *Or67d,* and *fru^M^* mutants. Instead, we noticed other *ppk* genes such as *ppk6*,*7*,*13*,*14*,*15*,*19* are altered in different mutant conditions. Fru^M^ seems to have a bidirectional role in regulating *ppk* gene expression, where it activates the expression of a subset of *ppk* genes (*ppk7*,*13*,*14*,*15*) while repressing the expression of others (*ppk6* and *ppk25*). One way to reconcile these differences is that multiprotein PPK complexes composed of combinations of different PPK subunits, and the stoichiometric levels of each *ppk* transcript in a given neuron can determine channel function. For example, misexpression of *ppk23*, which normally is not expressed in the antennal ORNs, can interfere with PPK channel function by disrupting the existing functional complexes in a given neuron, or forming new PPK complexes, thus affecting physiological properties. Another possibility is that the transcriptional changes in *fru^Lex^/fru^/4-40^* mutant are an output for eliminating all *fru^M^* transcripts, thus masking individual effects of each *fru^M^* isoform, such as *fru^MA^, fru^MB^, or fru^MC^*. And finally, it is also possible that the slight upregulation of *ppk25* in *Or47b* and *fru^M^* mutants as well as large changes in *Or67d* mutants may be due to global *fru^M^* changes in the whole antennae, or through retrograde neuromodulatory signaling from the antennal lobe.

Antennal sensilla contain cell types other than ORNs, such as glia-like cells and support cells of sensillum, as well as epithelial cells. Since our transcription data is from the whole antennae, one possibility we cannot exclude is that differences in antennal gene expression in different genetic and social conditions are readouts from non-neuronal cells or other ORNs. Even though we anticipate the immediate effects of *Or67d* and *Or47b* mutants to happen in the ORNs expressing these two receptors, signals from ORNs can lead to secondary changes in gene expression in non-neuronal cells within the sensillum. This also brings to light a general issue with bulk tissue where large cell-type-specific changes may be masked by cell-nonautonomous changes in gene expression in others cell types, as well as retrograde feedback signaling within olfactory circuits. Regardless, our data shows many of the differentially expressed genes encode regulators of neuronal function and neuromodulation. This increases the likelihood that the transcriptional changes in response to social and pheromonal cues are happening mostly in the neurons that respond to social cues, such as Or47b and Or67d ORNs. Future single-cell chromatin and transcription profiles from Fru^M^-positive neurons in the antenna and brain will provide deeper insights to neuron-specific changes in gene regulation from the peripheral to the central nervous system that modulate circuit function in response to social cues.

Part of the transcriptional effects can also be exerted downstream of changes in *dsx* levels seen in pheromone receptor and *fru* mutants. Given the upregulation of *dsx* levels in pheromone receptor and *fru* mutants suggests the possibility that some of the social experience-and neural activity-dependent transcriptional changes might also arise from increased Dsx. Dsx expression is restricted to non-neuronal cells in the antenna [58]. Similarly, neuromodulatory genes such as *Obps* and some neurotransmitter receptors, which function to alter both spontaneous and evoked activity of ORNs, are also expressed in non-neuronal cells in addition to the ORNs in the antennae [82–84, 88]. The social experience and pheromone signaling-dependent misregulation of these genes, points to adaptive homeostatic mechanism within local sensilla that can contribute to modulation of neuronal activity.

Lastly, in addition to the transcriptional changes occurring in neuromodulatory programs, genes regulating juvenile hormone metabolism are also modified with social context and pheromone receptor and *fruitless* mutants. Social experience works together with juvenile hormone signaling to modulate responses of pheromone sensing neurons in a Fru^M^ dependent manner [4]. These contribute to modification of competitive copulation advantage of males in different population densities and different ages as well as regulating overall courtship. At the molecular level social/pheromonal cues work together with juvenile hormone receptors to modulate chromatin around *fruitless* P1 promoter and its transcription [24]. Juvenile hormone acts as a repressor of *fru* expression and social experience converts it to an activator. In the same study, we showed that social isolation and disruption of Or47b signaling increases the accumulation of juvenile hormone receptor at the *fru* P1 promoter and juvenile hormone response elements.

This might be due to changing levels of juvenile hormone since our results show that expression of genes involved in juvenile hormone metabolism are altered in social isolation, and mutants in pheromone receptors and *fru*. The findings in our study together with results from previous studies suggest the presence of interconnected gene regulatory networks among social/pheromone signaling, hormone signaling and Fru^M^ function in neural and behavioral modulation (Figure 9).

## Conclusions

Social isolation is known to affect a wide range of brain functions and behaviors, such as aggression, attention, depression, and anxiety. Overall, this study highlights the shared transcriptional changes in master behavioral regulators and their target neuromodulatory genes providing a molecular mechanism that alter neural responses with social experience and pheromone sensing.

## Material and Methods

### Fly genetics and genotypes

Flies were raised on standard fly food (containing yeast, cornmeal, agar, and molasses) at 25°C in a 12-hour light/12-hour dark cycle in cylindrical plastic vials (diameter, 24 mm and height, 94 mm). For social isolation (single housing, SH) condition, 80-100 hour-old pupae were separated by sex and males were placed into individual vials, allowed to eclose alone, and aged to 7 days to deprive flies of pheromone interaction on ORNs. For group housing (GH) condition, 25-30 newly eclosed males were collected and placed into food vials. These were aged to 7 days and 180 antennae were dissected per sample, for a total of 3 samples for *w^1118^ GH*, 3 samples for *w^1118^ SH*, 3 samples for *Or47b^1^* mutants (*Or47b^1^/Or47b^1^; Or47b-GAL4, UAS-mCD8GFP/ Or47b-GAL4, UAS-mCD8GFP*), 3 samples for *Or67d^GAL4^* mutants (*UAS-mCD8GFP/ UAS-mCD8GFP; Or67d^GAL4^/Or67d^GAL4^*), and 2 samples for *fru^LexA^/ fru^4-40^* mutants (*w^+^; +/+; fru^LexA^/ fru^4-40^*).

### RNA-seq

RNA-seq was performed as described before (31). Male flies are aged for 7 days and dissected for the third antennal segment (∼180 antennae per genotype). RNA was extracted from dissected tissues samples using Qiagen RNA-easy extraction kit, quantified using a Qubit RNA assay kit and checked for quality using a High Sensitivity RNA ScreenTape on a TapesStation (Agilent). RNA integrity scores are typically 7.0 and greater. 1ug of RNA was used to construct libraries for sequencing using a KAPA mRNA library prep kit with polyA RNA selection. Barcoded libraries are sequenced on a Novaseq 6000 SP 50 bp following manufacturer’s instructions (Illumina). After demultiplexing sequence quality was assessed using FASTQC (Version 0.11.9). While there are issues with under-clustering of the samples and unbalanced pools, the data quality was typical for RNA extracted from fresh frozen material. The unbalanced pools resulted in differences in sequencing depth of each sample.

### Analysis of RNA-seq data

Once sequenced, the reads are preprocessed with FASTP [108] to remove adaptors and trim/filter for quality. These are mapped to the dm6 reference genome using MapSplice2 [109], with individual mapping rates exceeding 98% in all cases. This raw alignment was deduplicated and filtered for mapping quality and correct pairing; additional alignments are generated to confirm results are robust to mapping ambiguity. Mapped reads are assigned to genes in the annotation using the feature Counts command from the SubRead package [110]. Differential expression was modeled using DESeq2 [111] using the “apeglm” shrinkage estimator, and data was processed and visualized in R using the tidyverse framework [112], supplemented with the biomaRt [113], ComplexHeatmap [114] and UpSet [115] packages. The bioinformatics pipeline was implemented in Snakemake [116]. Code for the analysis is deposited on GitHub (https://github.com/csoeder/VolkanLab_BehaviorGenetics/tree/master/scripts). DEXSeq was used to test for differential exon use under models corresponding to those used in differential gene expression [46]. From the genome-wide test, the *fruitless* locus was examined in particular.

### Statistical analysis

Adjusted p-value were directly calculated from DESeq2 or DEXSeq (Supplementary table 1). Other statistical analysis is described in the legend of corresponding figures.

Specifically, to compare the exon usage in Figure 4, we also calculated p-value from post-hoc t-tests from raw read counts of independent comparisons of group-housed male antennae to each experimental condition at an individual exon segment (1-22, see the table 2 below). Even though many exons level differences were significant using this method, adjusted p-value from DEXSeq gave rise to fewer significantly altered exon levels.

**Table 1.** Genotypes used in in-vivo validation of expression.

| Figure | Type | Genotype | Source |
| --- | --- | --- | --- |
| Figure 6G,G' | Control | *+/+; 5-HT2A[2048-GAL4]/40XUAS-mCD8GFP* | *5-HT2A[2048-GAL4]*:<br>BDSC 66185 |
|  | *Or47b<sup>1</sup>* | *Or47b<sup>1</sup>/ Or47b<sup>1</sup>; 5-HT2A[2048-GAL4]/40XUAS-mCD8GFP* |  |
| Figure 6H,H' | Control | *dmGlut[0546-GAL4]/UAS-mCD8GFP; +/+* | *dmGlut[0546-GAL4]*:<br>BDSC 63397.<br>*Or67d<sup>Z3-5499</sup>*: Dean<br>Smith Lab, UT<br>Southwestern |
|  | *Or67d<sup>Z3-5499</sup>* | *dmGlut[0546-GAL4]/UAS-mCD8GFP;*<br>*Or67d<sup>Z3-5499</sup>/Or67d<sup>Z3-5499</sup>* |  |
|  | *fru<sup>LexA</sup>/fru<sup>4-40</sup>* | *dmGlut[0546-GAL4]/UAS-mCD8GFP;*<br>*fru<sup>LexA</sup>/fru<sup>4-40</sup>* |  |
| Figure 8D,D' | Control | *Jheh3[0892-GAL4]/UAS-mCD8GFP; +/+* | *Jheh3[0892-GAL4]*:<br>BDSC 63877.<br>*Or67d<sup>Z3-5499</sup>*: Dean<br>Smith Lab, UT<br>Southwestern |
|  | *Or67d<sup>Z3-5499</sup>* | *Jheh3[0892-GAL4]/UAS-mCD8GFP;*<br>*Or67d<sup>Z3-5499</sup>/Or67d<sup>Z3-5499</sup>* |  |
|  | *fru<sup>LexA</sup>/fru<sup>4-40</sup>* | *Jheh3[0892-GAL4]/UAS-mCD8GFP;*<br>*fru<sup>LexA</sup>/fru<sup>4-40</sup>* |  |
|  | *Or47b<sup>1</sup>/+* | *Or47b<sup>1</sup> Jheh3[0892-GAL4]/+; 40XUAS-mCD8GFP/+* |  |
|  | *Or47b<sup>1</sup>* | *Or47b<sup>1</sup> Jheh3[0892-GAL4]/Or47b<sup>1</sup>;*<br>*40XUAS-mCD8GFP/+* |  |

**Table 2.** t-test p-value of *fru* exons from DEXseq.

| t-test p-value | *w*<sup>1118</sup> *GH* vs |  |  |  |
| --- | --- | --- | --- | --- |
| Exon | *w*<sup>1118</sup> *SH* | *Or47b*<sup>1</sup> | *Or67d*<sup>GAL4</sup> | *fru*<sup>LexA</sup>/*fru*<sup>4-40</sup> |
| P1 (1) | 0.5696 | 0.0316 | 0.0619 | 0.0764 |
| Male (2) | 0.6843 | 0.0292 | 0.0125 | 0.0013 |
| Female (3) | 0.0697 | 0.3486 | 0.6932 | 0.5993 |
| P2 (4) | Expression too low to calculate |  |  |  |
| P6 (5) | Expression too low to calculate |  |  |  |
| P3 (6) | Expression too low to calculate |  |  |  |
| Exon 7 (7) | Expression too low to calculate |  |  |  |
| Exon 8 (8) | Expression too low to calculate |  |  |  |
| PD (9) | Expression too low to calculate |  |  |  |
| P4 (10) | 0.8788 | 0.0387 | 0.3002 | 0.1493 |
| P5 (11) | Expression too low to calculate |  |  |  |
| C1 (12) | 0.5489 | 0.0117 | 0.1050 | 0.0498 |
| C2 (13) | 0.6455 | 0.0826 | 0.0007 | 0.0079 |
| C3 (14) | 0.3622 | 0.1247 | 0.0399 | 0.0650 |
| C4 (15) | 0.8295 | 0.3811 | 0.0704 | 0.1205 |
| D (16) | 0.7090 | 0.2796 | 0.2787 | 0.7042 |
| C5 (17) | 0.7575 | 0.7338 | 0.2378 | 0.3174 |
| 3'UTR (18) | 0.7235 | 0.0086 | 0.0754 | 0.1328 |
| FruA (19) | 0.5241 | 0.0498 | 0.0171 | 0.0878 |
| FruB (20) | 0.8142 | 0.4809 | 0.7873 | 0.2550 |
| FruMC male (21) | 0.8519 | 0.1819 | 0.1969 | 0.1943 |
| FruFC female (22) | 0.4168 | 0.1123 | 0.8214 | 0.0223 |

### Quantitative reverse transcription PCR (qRT-PCR)

The qRT-PCR protocol was modified based on the previous protocol of the Volkan lab [32]. For each genotype (same as RNA-seq), four biological replicates were prepared separately, with each replicate containing 100 antennae from 50 males (7-day old). Antennae were dissected on Flypad and transferred into TRIzol (Invitrogen, 15596026) immediately. Total antennae RNA was extracted using the RNeasy Mini Kit (QIAGEN, 74104) and treated with Dnase I (TURBO DNA-free Kit, Invitrogen, Thermo Fisher Scientific AM1907) to remove genome DNA. cDNA was generated from the reverse transcription of 80-150ng total RNA using the SuperScript IV First-Strand Synthesis Kit (Invitrogen, 18091050) and poly d(T) as transcription primers. qPCR was performed using the FastStart Essential DNA Green Master kit (Roche, 06924204001) on LightCycler® 96 instrument (Roche, 05815916001). Primers used are listed in Table 3 below. The expression level was calculated by ΔCt method using the *fl(2)d* as the standard gene. The calculation was performed in GraphPad Prism software. One-way ANOVA was used for significance test, followed by multiple comparisons (compare other groups to group-housed wild types *w^1118^ GH*). * p<0.05, ** p<0.01, *** p<0.001, **** p<0.0001.

**Table 3.**
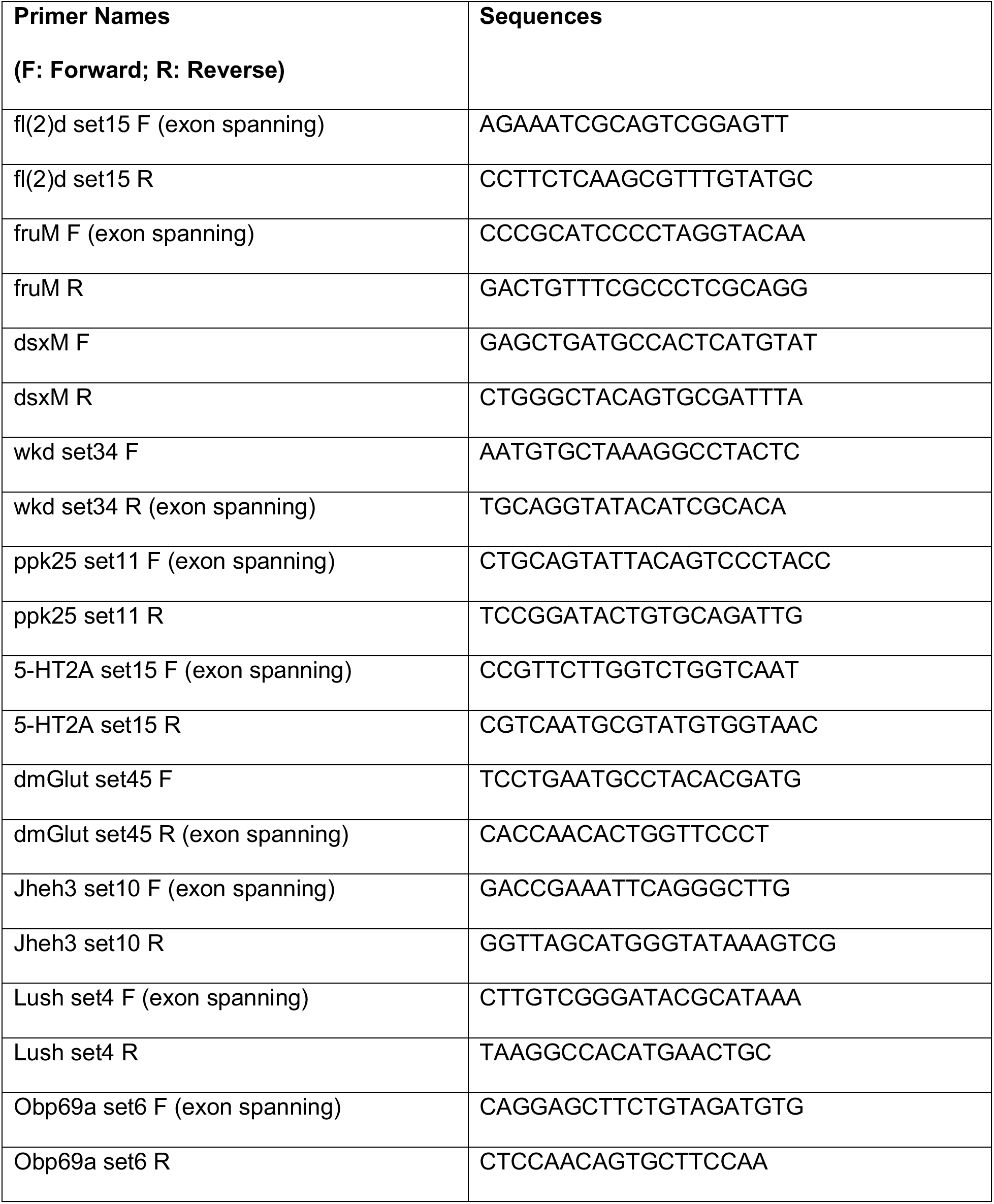
Primers sets used in qRT-PCR assays.

### In-vivo validation of gene expression

Fly heads were dissected in cold PBT (phosphate buffered saline with Triton X-100) buffer and were fixed in 4% PFA on nutator at room temperature for 1 hour. Fly heads were washed three times with fresh PBT, and every wash was 10 min at room temperature. Antennae were dissected from heads and were fixed in 4% PFA on nutator at room temperature for 30 minutes. Antennae were washed three times with fresh PBT, and every wash was 10 min at room temperature. Antennae were mounted using Fluoromount-G Slide Mounting Medium (SouthernBiotech). Images were taken by Olympus FluoView FV1000 confocal microscope. Controls and experimental groups were imaged by same parameters. Native fluorescence was measured by ImageJ. The fluorescence intensity was defined as the fluorescence of region of interest subtracted by that of the background. Statistical tests were performed in GraphPad Prism software. One-way ANOVA was used for significance test, followed by multiple comparisons (compare other groups to group-housed control). * p<0.05, ** p<0.01, *** p<0.001, **** p<0.0001.

## Acknowledgements

We are grateful to Liqun Luo and Hongjie Li for sharing the single-cell RNA-seq data from adult ORNs prior to publication. We would like to thank Yetong Huang, George Thomas Barlow, and Paulina Guerra-Schleske for their contributions to sample collection and RNA extraction of samples, and the Volkan lab for help with the manuscript. We thank the Bloomington Stock Center and UNC High Throughput Sequencing Facility for their services.

## Contributions

Conceptualization/design of the work: BD and PCV. Investigation: CD, QD, and BD with help from AM and DG. Analysis/interpretation of data: QD, BD, CD, CS, CDJ, and PCV. Drafted the work: BD, QD, CD and PCV. The authors read and approved the final manuscript.

## Ethics declarations

### Consent for publication

Not applicable.

## Availability of data and materials

All relevant data are within the paper and its supporting information files. Raw sequencing data will be uploaded to the GEO under embargo pending the submission and acceptance of the data submitted here for publication.

---

**Figure 1-figure supplement 1.**
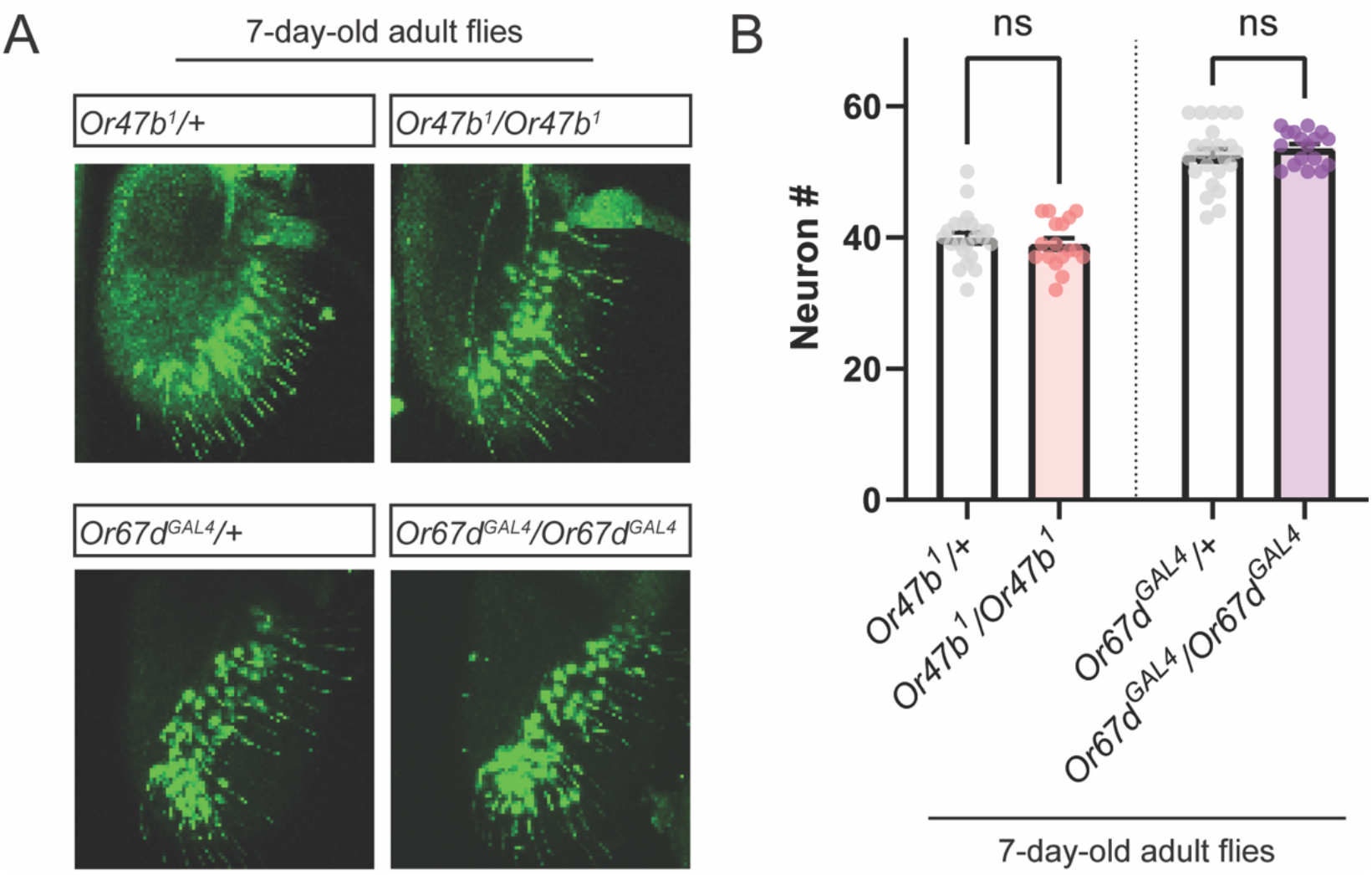
ORN numbers of control and mutants. **(A)** *<u>Or47b-GAL4</u>* driven *UAS-mCD8GFP* expression in 7-day old male antennae from control and *Or47b* mutants (top), and *Or67d^GAL4^* driven *UAS-mCD8GFP* expression in 7-day old male antennae from control and *Or67d* mutants. **(B)** Statistics of numbers of Or47b and Or67d ORNs in both control and *Or* mutants.

**Figure 4-figure supplement 1.**
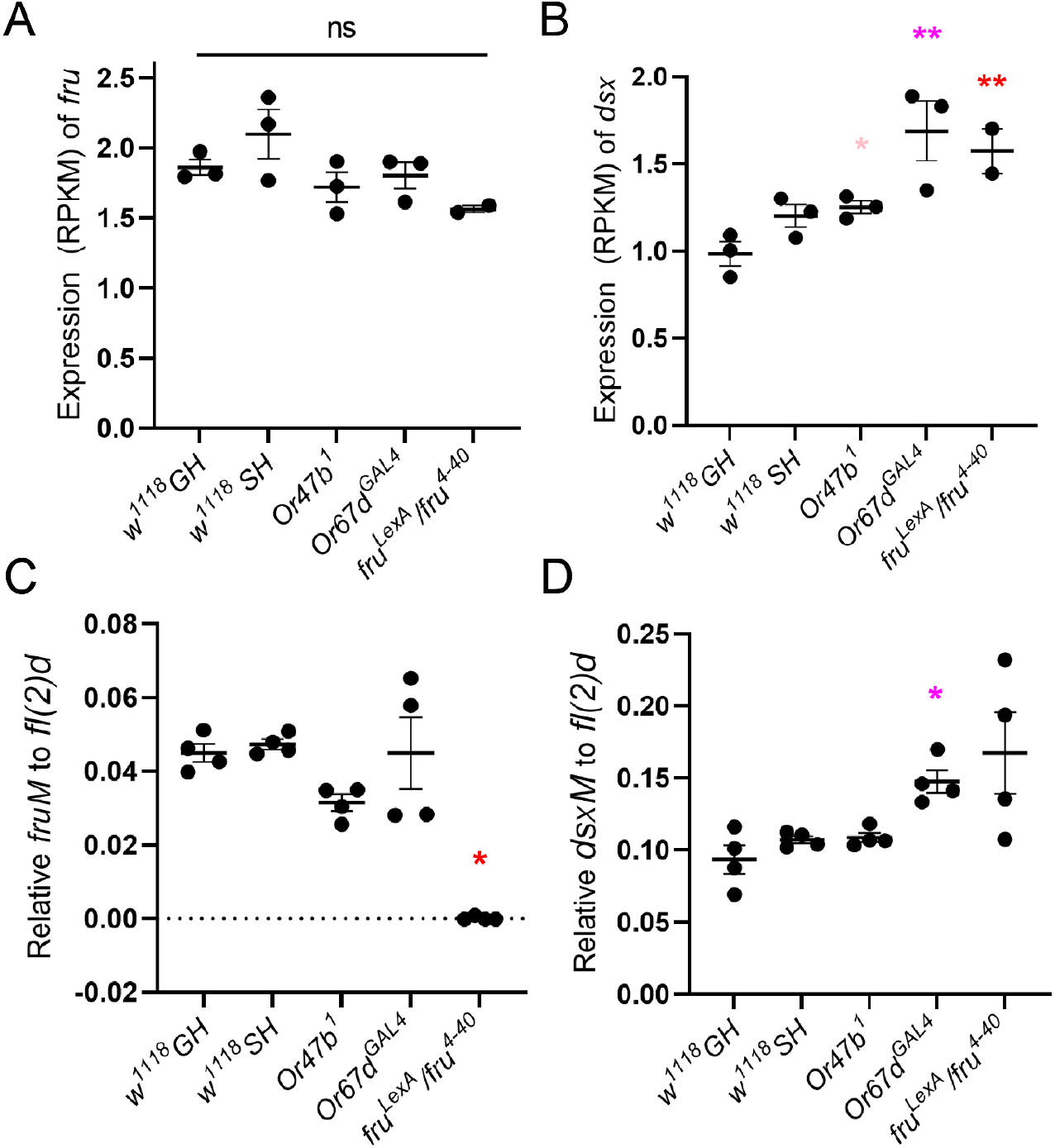
Validation of *fru* and *dsx* male specific exon expression in male antennae of WT and mutants. **(A-B)** RPKM of *fru* and *dsx.* Adjusted p-value was directly performed via DESeq2. **(C-D)** Quantitative RT-PCR validation of *fru* and *dsx* male specific exon expression. **(C)** The expression of *fru^M^* was not detected in *fru* mutants and did not show significant difference in other *Or* mutants or SH WT. **(D)** The *dsx^M^* expression showed significant increase in *Or67d* mutant male antennae.

**Figure 5-figure supplement 1.**
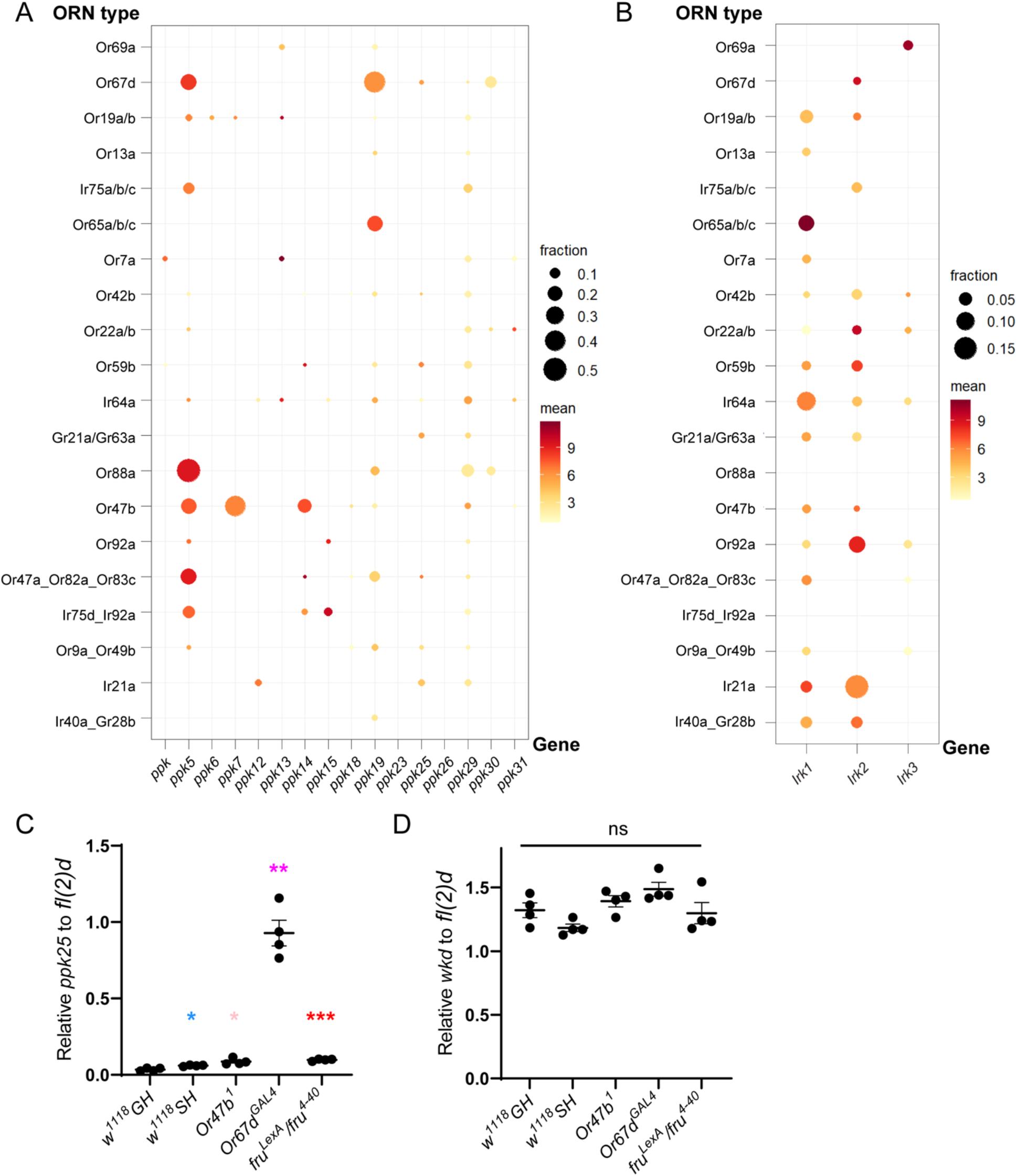
Validation of *ppk* expression across ORN classes and in mutants. **(A-B)** ORN class-specific expression of *ppk* and *Irk* family genes based on single-cell RNA-seq datasets from the adult ORNs [66]. Size of each circle indicates the fraction of positive cells (log_2_(CPM+1)>0.5) and color intensity indicates the mean expression (log_2_(CPM+1)) of all positive cells. **(C-D)** Quantitative RT-PCR validation of *ppk25* expression (C) and a negative control gene *wkd* expression (D) from antennae of grouped and socially isolated wild types, *Or47b* mutants, *Or67d* mutants, and *fru^M^*mutants normalized to *fl(2)d*. *fl(2)d* and *wkd* were selected based on their near-identical expression level across all conditions from the RNA-seq results and thus used as loading and negative control genes.

**Figure 6-figure supplement 1.**
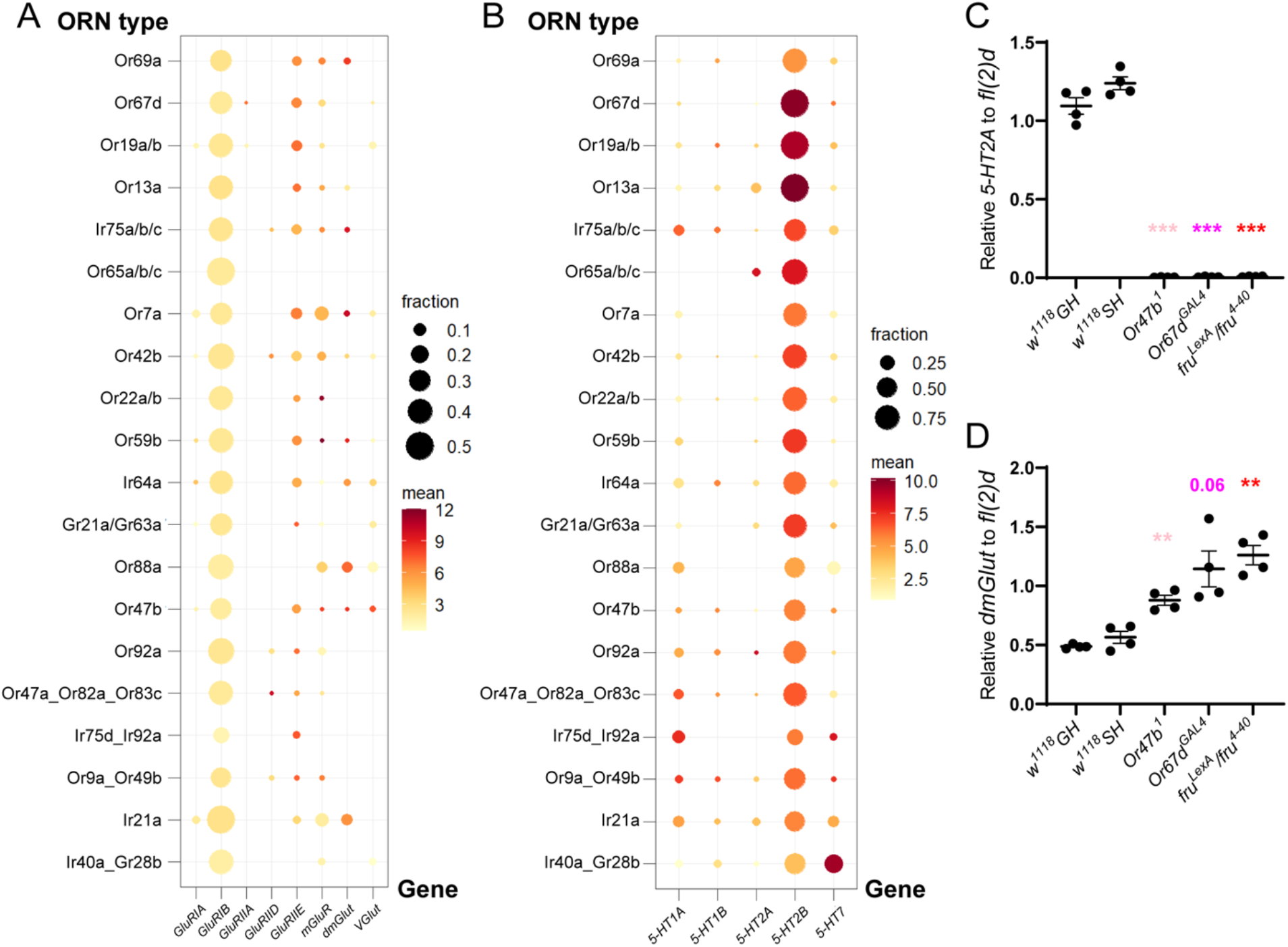
Validation of neurotransmitter receptor expression across ORN classes and in mutants. **(A-B)** ORN class-specific expression of serotonin and glutamate receptors based on single-cell RNA-seq datasets from the adult ORNs [66]. The size of each circle indicates the fraction of positive cells (log_2_(CPM+1)>0.5) and color intensity indicates the mean expression (log_2_(CPM+1)) of all positive cells. **(C-D)** Quantitative RT-PCR validation of *5-HT2A* **(C)** and *dmGlut* **(D)** expression from antennae of grouped and socially isolated wild types, *Or47b* mutants, *Or67d* mutants, and *fru^M^* mutants normalized to *fl(2)d*.

**Figure 7-figure supplement 1.**
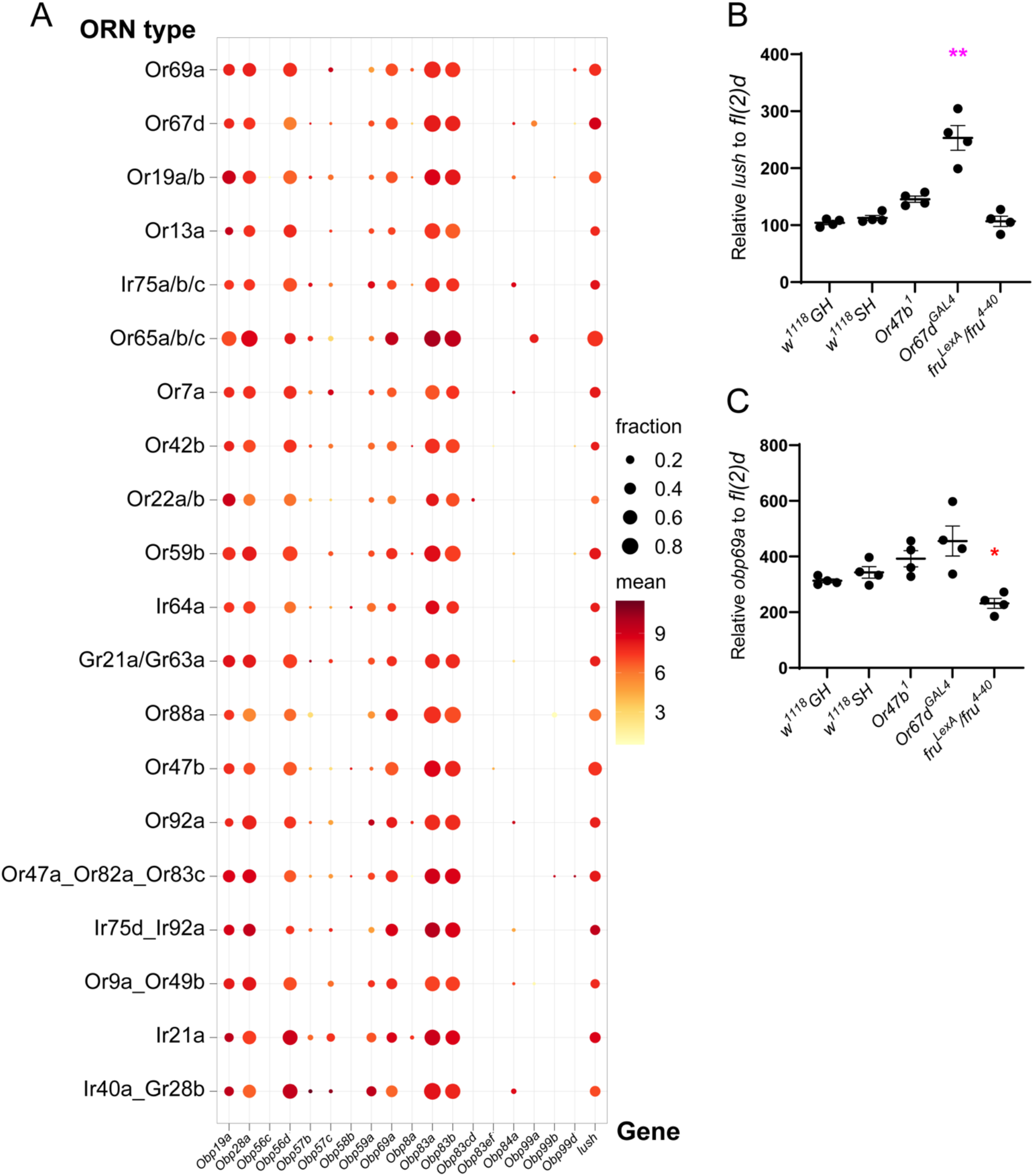
Validation of *Obp* expression across ORN classes and in mutants. **(A)** ORN class-specific expression of *Obps* based on single-cell RNA-seq datasets from the adult ORNs [66]. The size of each circle indicates the fraction of positive cells (log_2_(CPM+1)>0.5) and color intensity indicates the mean expression (log_2_(CPM+1)) of all positive cells. **(B-C)** Quantitative RT-PCR validation of *lush* **(B)** and *Obp69a* **(C)** expression from antennae of grouped and socially isolated wild types, *Or47b* mutants, *Or67d* mutants, and *fru^M^* mutants normalized to *fl(2)d*.

**Figure 8--figure supplement 1.**
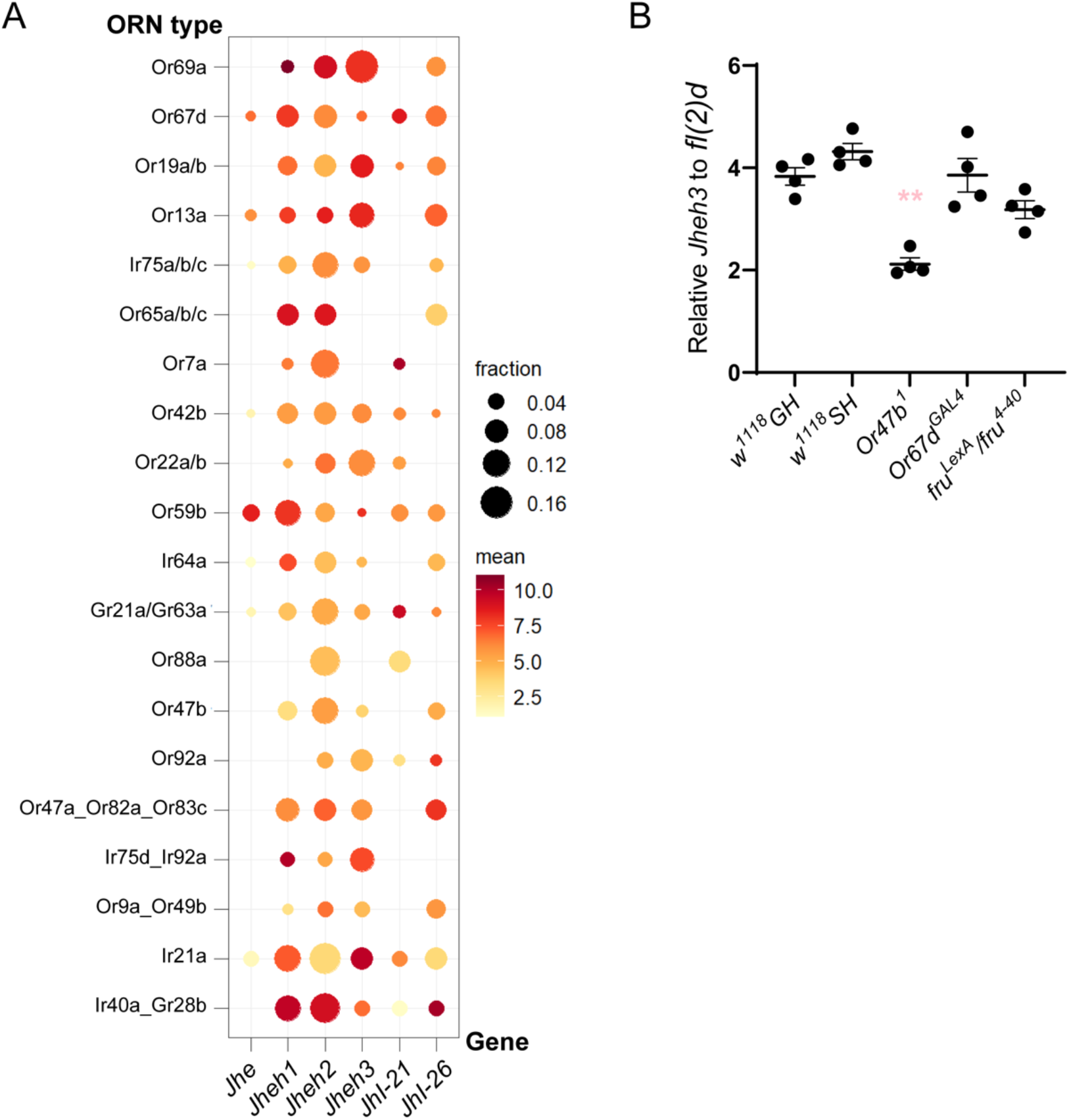
Validation of hormone-related gene expression across ORN classes and in mutants. **(A)** ORN class-specific expression of juvenile hormone regulators based on single-cell RNA-seq datasets from the adult ORNs [66]. The size of each circle indicates the fraction of positive cells (log_2_(CPM+1)>0.5) and color intensity indicates the mean expression (log_2_(CPM+1)) of all positive cells. **(B)** Quantitative RT-PCR validation of *Jheh3* expression. Relative expression of *Jheh3* from antennae of grouped and socially isolated wild types, *Or47b* mutants, *Or67d* mutants, and *fru^M^* mutants normalized to *fl(2)d*.

